# A morphological basis for path dependent evolution of visual systems

**DOI:** 10.1101/2022.12.20.520810

**Authors:** Rebecca M. Varney, Daniel I. Speiser, Johanna Cannon, Morris Aguilar, Douglas J. Eernissee, Todd H. Oakley

## Abstract

Path dependence influences macroevolutionary predictability by constraining potential outcomes after stochastic evolutionary events. Although demonstrated in laboratory experiments, the basis of path dependence is difficult to demonstrate in natural systems because of a lack of independent replicates. Here we show two types of complex distributed visual systems each recently evolved twice within chiton mollusks, demonstrating rapid and path dependent evolution. The type of visual system a chiton lineage evolves is constrained by the number of openings for optic nerves in its shell plates: lineages with more openings evolve visual systems with thousands of eyespots, whereas those with fewer evolve visual systems with hundreds of shell eyes. These macroevolutionary outcomes shaped by path dependence are both deterministic and stochastic because possibilities are restricted yet not entirely predictable.

**One-Sentence Summary:** Multiple convergent origins of visual systems show macroevolution of complex traits can be rapid and contingent upon pre-existing structures

## Main Text

Establishing the extent to which the evolutionary trajectories of complex systems are contingent on historical events - a phenomenon termed path dependence (*1*, *2*) - is fundamental for understanding the predictability of evolution. If evolution is predictable, single, locally optimal solutions to environmental problems will tend to emerge; if stochastic, many, functionally equivalent solutions will solve the same problem (*3–6*). Path dependence occurs when evolutionary paths contain “critical junctions”, which we define as stochastic events that split evolutionary trajectories so that lineages commit to one of multiple possible evolutionary pathways, thereby constraining the suite of possible outcomes. Path dependence is well established for the evolution of particular proteins and in unicellular laboratory systems (e.g. (*7–13*)). Furthermore, some specific evolutionary outcomes are known to be restricted by earlier stochastic events in a few natural systems (e.g. *14–16*).

Demonstrating path dependence in natural systems is challenging because it requires the identification of critical junctions and elucidation of the constraints those junctions impose on future evolutionary paths (*17–20*). First, critical junctions are difficult to identify because alternative evolutionary pathways are often not observable along the singular history of life. Path dependence may be inferred from convergent evolution because convergent origins illustrate multiple evolutionary pathways, analogous to replicates in controlled laboratory experiments. If splits in evolutionary trajectories lead to functionally equivalent outcomes in convergent lineages, those splits could be identified as critical junctions. Second, even if convergent evolution reveals potential critical junctions, convergent evolution occurs most commonly in traits of organisms with very different body plans and ecologies. Different organismal and environmental contexts are likely to exert different selective pressures on traits, so the majority of instances of convergent evolution are not effective replicates for establishing path dependence (*12*). Finally, accurately reconstructing the evolutionary histories of convergent traits in multiple taxa requires understanding of ancestral conditions, as well as knowledge of the timing of key transitions in character states. This requires a detailed fossil record and a robust phylogenetic history beyond that which is available for many taxa. Together, these obstacles make identification of critical junctions and path dependence in natural systems enormously challenging.

By overcoming the challenges imposed by other natural traits, the visual systems of chitons (Mollusca; Polyplacophora) provide a compelling case to test hypotheses about path dependent evolution. First, morphologically distinct visual systems appear to have evolved in separate lineages of chitons. Chiton visual systems likely evolved from aesthetes, which are numerous, microscopic sensory organs embedded in the shell plates (*21*). In some lineages, eyespots (20-35 μm wide) are attached to aesthetes (*22*, *23*). In other lineages, the aesthetes are interspersed with camera-type eyes with image-forming lenses made of shell material (up to 145μm wide), hereafter referred to as shell eyes (*21*, *24–27*). Both the eyespot- and shell eye-based distributed visual systems of chitons provide spatial vision (*26–28*). If eyespots and shell eyes evolved separately in chitons, these distributed visual systems may represent distinct evolutionary paths to a convergent functional outcome: spatial vision. Second, chitons have a relatively rich fossil record, permitting time-calibrated phylogenetic analyses. If the distributed visual systems of chitons have recent origins, their evolutionary histories may be reconstructed with greater confidence than those of other visual systems, which largely have ancient histories (*29*). Finally, fossil and extant chitons are found in similar environments, and so tend to be ecologically similar: most species live (or lived) on hard substrates in intertidal or shallow subtidal habitats. The body plan present in both fossil and extant chitons is consistent across clades and evolutionary time (*30–33*). Species exhibiting the full range of shell-embedded sensory organs can even be found living on the same rock (*34*). Here, we ask if the evolution of distributed visual systems in chitons is path dependent by mapping the origins of eyespots and shell eyes onto the most comprehensive chiton phylogeny produced to date. We then use the rich fossil record of chitons to time-calibrate our phylogeny and identify specific points where stochastic events committed some lineages to one or another specific evolutionary pathway, resulting in multiple solutions to the evolution of visual systems. Our discoveries highlight that the evolution of complex visual systems, often portrayed as deterministic (e.g. (*35*)), is path dependent: stochastic events at specific points - critical junctions - constrain lineages to one of a subset of relatively determined pathways.

### Phylogenomics shows chitons rapidly evolved visual systems four times in two distinct forms

To characterize patterns of visual system evolution in chitons, we used genomic target capture and Bayesian inference to produce the most complete phylogeny of chitons to date, with emphasis on Chitonina, the suborder that includes over half of all extant chiton species and most chitons with eyespots or shell eyes. We discovered distributed visual systems evolved separately in chitons at least four times: two lineages via eyespots and two other lineages via shell eyes (Figure 1A/B). The two lineages that contain species with eyespots, Callochitonida (*23*) and Chitonidae:Chitoninae (*22*, *24*), are distantly related to one another, consistent with previous molecular phylogenies (*36–38*). Likewise, distributed visual systems based on shell eyes also evolved twice separately: once in Chitonidae: Acanthochitoninae+Toniciinae and once in *Schizochiton*, which in our phylogeny falls as sister to the rest of Chitonina, making it a distant relative of Acanthopleurinae+Toniciinae. The placement of *Schizochiton* has been uncertain across studies of chitons (*36*, *39*, *40*), in part because Schizochitonidae contains only two accepted species, *Schizochiton incisus* (*39*) and *Schizochiton jousseaumei* (Dupuis, 1917), and *S. incisus* are the only specimens available. However, we are confident in our results: tests of branch stability do not indicate phylogenetic uncertainty in the placement of *S. incisus* in any of our analyses (Supplementary Material, Leaf Instability Testing).

**Fig. 1:**
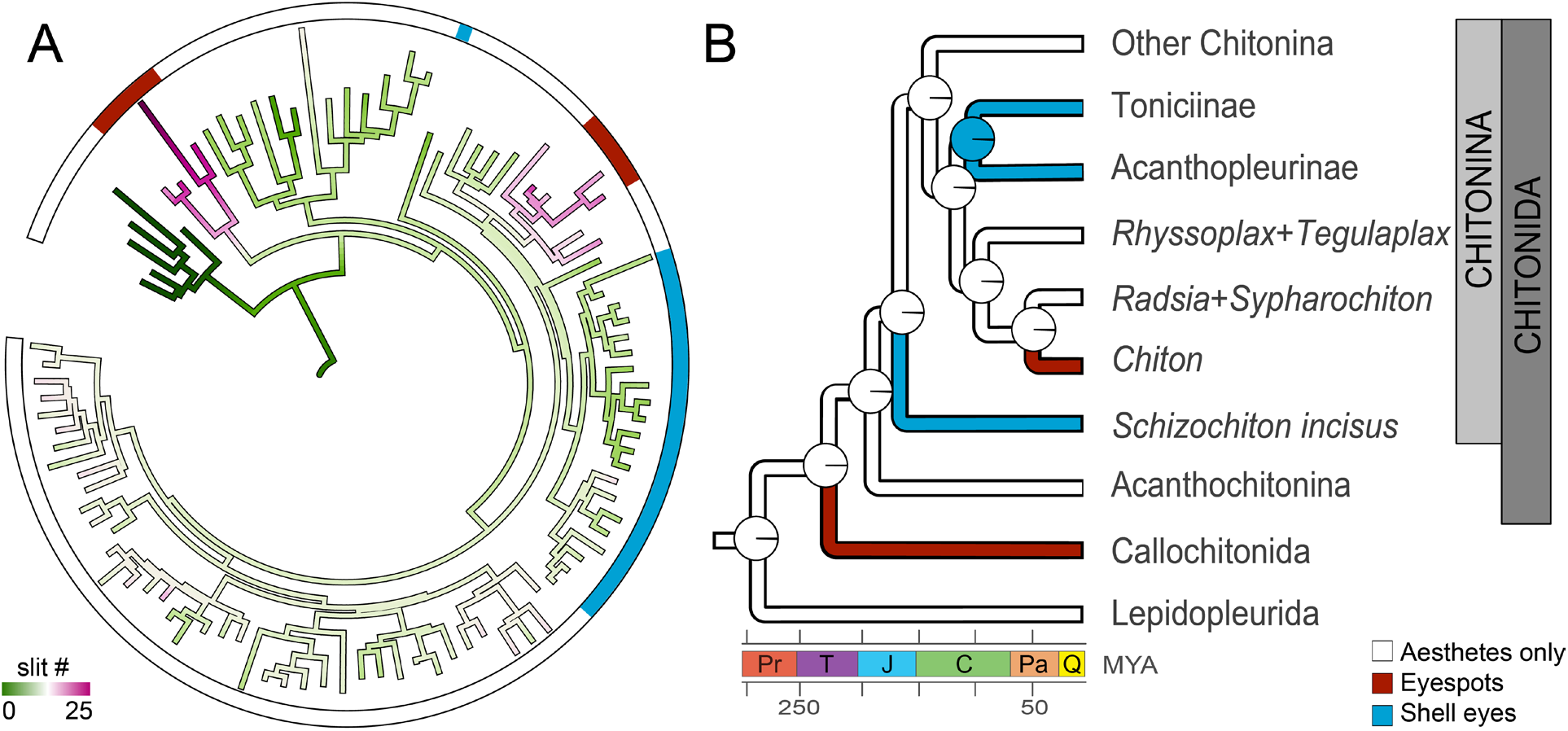
Two types of distributed visual systems evolved convergently in chitons, one type based on eyespots (red) and the other on shell eyes (blue). **(A)** The full maximum likelihood phylogeny of chitons produced by this study (outgroups not shown). Branch coloration indicates ancestral state reconstruction (ASR) of the number of slits in the anteriormost shell plate, where dark green represents 0 slits and pink represents >10 slits. **(B)** A time-calibrated phylogeny generated with Bayesian inference, showing four independent origins of visual systems in chitons. Divergence times correspond to the geologic time scale below. ASR implies that all origins of eyespots or shell eyes in chitons come from an aesthete-only starting point, where all proportional marginal likelihoods are >95%. Additional support metrics are included in the supplemental materials.

Next, to assess support for independent origins of distributed visual systems in chitons, we performed ancestral state reconstruction (ASR). Using ASR, we found high support (≥95% proportional marginal likelihood) for all four instances of visual system evolution in chitons occurring independently. Not only all Chitonina but all living chitons and even ancient fossil polyplacophorans (*32*) have or had aesthetes but, as ASR reveals, eyespots and shell eyes are surprisingly recent and non-homologous components of chiton visual systems. Eyespots evolved independently in Callochitonida and Chitoninae, and likewise shell eyes evolved independently within Chitonidae and in Schizochiton. Within Chitonidae, eyespots evolved in a subclade in Chitoninae and shell eyes evolved in the last common ancestor of the other two subfamilies, Acanthopleurinae and Toniciinae. ASR indicates these visual systems evolved separately: the last common ancestor of Chitonidae only had aesthetes (95% proportional marginal likelihood). Each of the four separate origins of visual systems is inferred to have evolved independently in chitons, including convergent origins of two different types of distributed visual systems, one based on eyespots and the other on shell eyes (Figure 1A). Further, we find eyespots and shell eyes have evolved from aesthetes separately, rather than eyes evolving from eyespots that evolved from aesthetes, a stepwise pattern that would have corresponded to the relative levels of morphological complexity in these organs.

To understand the timing of separate origins of visual systems in chitons, we used fossil occurrence data to time-calibrate the phylogeny and found all four convergent visual systems in chitons are less than 200 million years old (Figure 1B). Both instances of shell eyes in chitons represent the most recent origins of camera-type eyes known. By comparison, the more ancient camera-type eyes of vertebrates and cephalopods originated at least 500 and 425 mya, respectively (*29*, *41*). Distributed visual systems based on eyespots may have evolved even more recently than those based on shell eyes. To determine how rapidly chitons evolved visual systems based on eyespots, we quantified the time between the origin of eyespots in *Chiton* and the most recent eyeless ancestor in this clade. Using our fossil-calibrated time tree, we found that eyespots in *Chiton* originated within the last six million years. Theoretical models estimate eyes can evolve within 363,992 generations (*3*, *42*)), so if we assume chitons have generation times of three years (based on available studies of other chiton genera, e.g. (*43*)), a lineage could evolve a visual system in two million years, within an order of magnitude of the 6 million years we estimate for visual system evolution in *Chiton*. For comparison, the only published estimate of the time required to evolve an eye is from vertebrates, where eyes evolved in approximately 30 million years (*29*, *44*). Recent origins of shell eyes and eyespots in chitons allow us to calibrate the timing of visual system evolution with greater confidence from fossils, and the rapid acquisition of eyespots allows us to better understand the morphological changes associated with the origins of visual systems. Thus, the recent origins and rapid evolution of visual systems in chitons make them particularly valuable for understanding how complex traits evolve.

### Number of slits in shell plates indicate a critical junction in the evolution of chiton visual systems

To demonstrate path dependence in natural systems, we must identify critical junctions in the evolution of those systems. To discover critical junctions, we examined morphological differences between chitons with only aesthetes, chitons with eyespots, and chitons with shell eyes. Most chitons integrate their shell-embedded sensory organs, including their aesthetes, eyespots, and shell eyes, into their nervous system by passing nerves through slits along the edges of their shell plates (*21*, *39*, *45–47*). These slits are thus a vital part of chiton visual systems, as they make space for optic nerves to exit shell plates. We predicted that as chitons add sensory organs (like aesthetes, eyespots, and shell eyes), they would require larger or more numerous slits for additional nerves to pass through. We quantified the number of slits in the anteriormost shell plates of all chitons in our phylogeny (Supplemental Data). Members of the earliest branching clade of extant chitons, Lepidopleurida, lack slits. They innervate their aesthetes by running nerves through pores in their relatively thin shell plates, and outgroup comparisons indicate this is the ancestral state of crown-group chitons (*48*, *49*). In the remaining orders of chitons, Chitonida and Callochitonida, species have thicker shell plates, so slits are necessary to innervate aesthetes; all these chitons have at least eight or nine slits on their anterior shell plates (*31*). Both of the separate clades of chitons that have shell eyes retain the ancestral eight or nine slits. However, all chitons with eyespots have anterior shell plates with between 15 and 22 slits. We thus hypothesized that a higher number of slits is a critical junction in the evolution of visual systems in chitons, favoring the evolution of eyespots but not shell eyes.

If an increased number of slits is a critical junction that imposes a functional constraint favoring the evolution of eyespots instead of shell eyes, we would expect that an increase in the number of slits predates the origin of eyespots themselves. Within Chitoninae (Figure 1A), we compared the number of slits between species with eyespots and those with only aesthetes. All species with eyespots have ⋝15 slits across their anteriormost shell plates. *Chiton cumingsii*, sister to the remaining members of *Chiton* in our phylogeny, does not have eyespots but has 14 slits (personal observation by DE and DIS). Sister to *Chiton*, species in the *Radsia+Sypharochiton* clade have 13-16 slits. Sister to *Chiton+Radsia+Sypharochiton*, species in the *Rhyssoplax+Tegulaplax* clade have 9-10 slits. We performed ancestral state reconstruction and found that the ancestor of Chitonidae most likely had 8-9 slits, while the ancestor of Chitoninae likely had 10-11 slits (Supplementary Figure 3). Thus, slits became more numerous in Chitoninae prior to the evolution of eyespots in *Chiton*. In Callochitonida, the other clade of chitons that evolved eyespots, an eyeless sister species or clade is unavailable because all Callochitonida examined to date appear to have eyespots (*23*). All species of Callochitonida in our analysis have >15 slits. Ancestral state reconstruction suggests that the last common ancestor of Callochitonida and Chitonida had 8-9 slits, indicating an independent origin of the increased number of slits in Callochitonida. In contrast, chitons with shell eyes have 8 or 9 slits in their anterior shell plates, including *S. incisus* and the ancestor of Toniciinae+Acanthopleurinae. The evolution of eyespots in chitons appears to follow an increase in slit number, but the evolution of shell eyes does not.

### A phylomorphospace of chiton visual systems supports path dependent evolution

In light of the four separate origins of visual systems in chitons, and of the different morphological characters associated with them, we hypothesized that evolutionary outcomes in chiton visual systems were constrained by path dependent evolution. We further hypothesized that critical junctions have resulted in gaps in the morphospace of chiton visual systems due to the absence of intermediate forms: chitons can have eyespots or shell eyes but not visual systems that share morphological characters with both.

To test our hypotheses about path dependent evolution in chitons, we constructed a phylomorphospace based on morphological traits associated with visual systems: aesthete density and number of slits in anterior shell plates. We quantified aesthete density from available SEM images of chiton valves for species in our phylogeny and plotted it alongside the number of slits in the anteriormost valve of each species. Consistent with our prediction, we found a pronounced gap in the phylomorphospace that suggests slit number acts as a constraint on the type of visual system a lineage of chitons can evolve (Figure 2A, purple triangle). The absence of intermediates in the phylomorphospace of chiton visual systems shows that chitons evolved vision via one of two distinct paths and suggests that the morphological characters that define each visual system are mutually exclusive. Slit number appears to predict the evolution of eyespots in chitons. The number of slits in the anterior shell plates of chitons with eyespots is consistently higher than the ancestral number of slits, and only those lineages of chitons that increased the number of slits in their plates evolved eyespots. Slit number therefore acts as a critical junction, a stochastic change that commits a lineage of chitons to one or the other evolutionary pathway towards spatial vision. These critical junctions make the evolution of visual systems in chitons path dependent, visible as a gap in phylomorphospace created by mutually exclusive solutions for spatial vision that are morphologically divergent yet functionally convergent.

**Fig. 2.**
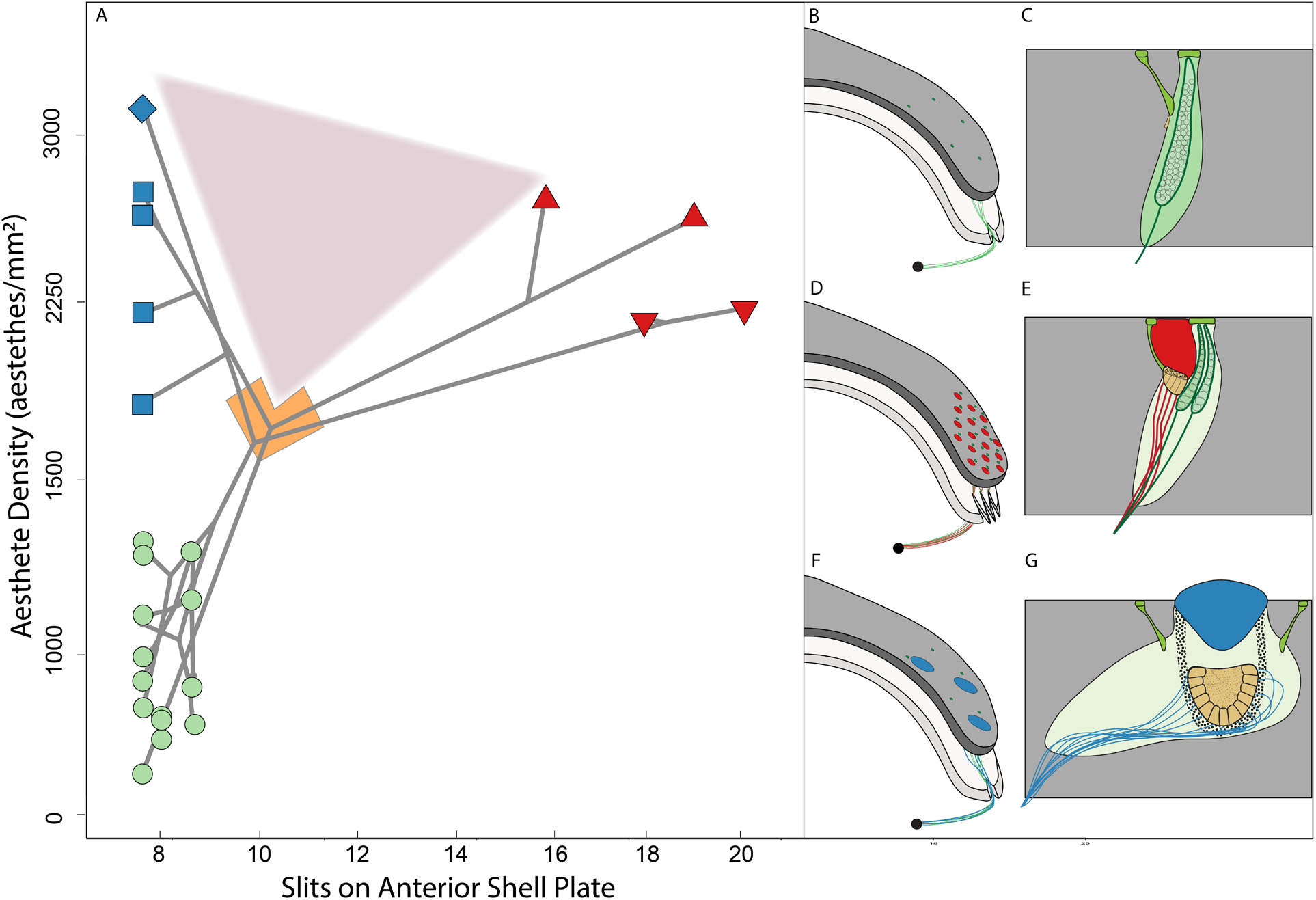
Path dependence of visual system evolution in chitons is indicated by a phylomorphospace of vision-related morphological traits, showcasing two mutually exclusive solutions to vision. **(A)** Phylomorphospace of chiton visual systems with number of slits on the anteriormost shell plate on the x-axis and aesthete density (in aesthetes/mm2) on the y-axis. There are two separate origins of shell eyes (blue squares and blue diamond) and two separate origins of eyespots (red triangles and red inverted triangles), both evolving from chitons with only aesthetes (green circles). We hypothesized that there would be a gap in the phylomorphospace, and we see such a gap (purple triangle). The gap results from the absence of intermediate forms of visual systems in chitons. Lineages are committed to one or the other visual system at a critical junction (orange polygon). **(B)** the distribution of aesthetes (green) across the surface of a shell plate. **(C)** the internal morphology of an aesthete (green), the simplest sensory structure embedded in the upper layer of the shell plate (grey). Micraesthetes (lime) branch off from macraesthetes (green) **(D)** the distribution of eyespots (red) and aesthetes (green) across the surface of a shell plate, and an illustration of multiple slits at the edge of the shell plate that facilitate the exit of the many nerves running through the shell plate. **(E)** the internal morphology of an aesthete with an attached eyespot, with a patch of photoreceptor cells (orange) beneath a clear portion of the shell plate (red) and a nerve running through the upper layer of the shell plate (grey). **(F)** the distribution of shell eyes (blue) and aesthetes (green) across the surface of a shell plate. **(G)** the internal morphology of a shell eye, with photoreceptor cells forming a retina (orange) beneath a lens (blue) and a large nerve running through the upper layer of the shell plate (grey). (B-G not to scale).

We found aesthete density varies between species across our phylogeny, but do not consider an increased in aesthete density to act as a critical junction. Chitons with eyespots and chitons with shell eyes both have a greater density of aesthetes than most other members of Chitonida (Figure 2A). Because all four lineages of chitons that gained visual systems share an increased density of aesthetes, we consider an increase in aesthete density to be a preadaptation for evolving a visual system. However, an elevated aesthete density does not appear to limit chitons to one or another type of visual system. When aesthete density increases but slit number does not, lineages evolve shell eyes but not eyespots.

## Conclusion

Convergent evolution is often portrayed as an inevitable feature of lineages moving toward an optimal solution to an environmental problem (*50*). Such arguments dismiss contingency as a lesser force than selective pressure, and assert that given sufficient time, an optimized trait will evolve in a deterministic manner. Yet chiton visual systems present a morphospace with multiple optima: networks of either eyespots or shell eyes provide chitons with spatial vision. We find evidence for a critical junction that defines convergent evolutionary pathways to visual systems in chitons, where lineages split into two discrete trajectories that led to mutually exclusive types of visual systems. A gap on a phylomorphospace of chiton visual systems suggests that chitons are constrained to a given evolutionary path because no intermediate visual systems appear. The two types of distributed visual systems of chitons rely on differing morphological innovations, like increased slit number in shell plates, that predate the evolution of spatial vision. Thus, ultimate evolutionary outcomes are constrained by earlier evolutionary events. Previous studies, confined to molecular biology in laboratory environments, demonstrated path dependence can dictate the order of adaptations and the persistence of changes across evolutionary time (*2*, *9*, *51*). Here, we demonstrate path dependence in a naturally evolving system. Evolution is as much historical as biological, so clarifying the role of history in shaping evolutionary outcomes is critical to our understanding of how and why characters can evolve in predictable ways.

**Extra Supp Refs:** (*11*, *23*, *28*, *30*, *31*, *36*, *46–49*, *52–92*)

## Supporting information

All Supplementary Materials

## Acknowledgments

We thank Philippe Bouchet, Barbara Buge and Nicolas Puillandre (MNHN, Paris), Gustav Paulay, John Slapcinsky and Amanda Bemis (UF, Florida Museum of Natural History), Daniel Geiger and Vanessa Delnavaz (SBMNH, Santa Barbara, CA), Terrance Gosliner and Christina Piotrowski (CASIZ, San Francisco, CA) for help with their respective collections. The authors are grateful to the following who provided specimens and aided us to obtain speciments in accordance with each country’s collection regulations: Irma Shita Arlyza, Paul Barber, Lesley Brooker, Carlos Cáceres Martínez, Bruno Dell’Angelo, Richard Emlet, António (Jose) França, Michel Hendrickx, Alan Hodgson, Christian Ibáñez, Mark Langdon, Susanne Lockhart, Peter Marko, Tomoyuki Nakano, Ron and Joy Noseworthy, Raphael Sagarin, Boris Sirenko, Julia Sigwart, H. William Detrich, Craig Starger, Shawn Wiedrick, Michele Weber, Demian Willette, and Craig Young. Authors thank Joanna Wolfe, Niko Hensley, and members of the Oakley lab for insightful remarks on the manuscript. JC thanks Andrew Swafford for assistance with code. We also thank Lesley Brooker and Kevin Kocot for providing additional chiton transcriptome data and support.

## Funding

National Science Foundation DEB 1354831 (DIS, THO)

National Science Foundation DEB 1355230 (DJE)

National Science Foundation EAGER 1045257 (THO)

National Science Foundation IOS 1754770 (THO)

## Author contributions

Conceptualization: DIS, DJE, THO

Methodology: RMV, DIS, JC, MA, DJE, THO

Investigation: RMV, DIS, JC, MA, DJE

Visualization: RMV

Funding acquisition: DIS, DJE, THO

Project administration: THO

Supervision: THO

Writing – original draft: RMV, DIS, THO

Writing – review & editing: RMV, DIS, JC, MA, DJE, THO

## Competing interests

Authors declare that they have no competing interests.

## Data and materials availability

All data and code used in the analyses are available in the main text, the supplementary materials, and in Dryad repository XXXX.

## Supplementary Materials

Please see included PDF.

## Notes

### Competing Interest Statement

The authors have declared no competing interest.

