## Supplementary material for "A morphological basis for path dependent evolution of visual systems": All Supplementary Materials

2022

**2 Supplementary Figures and Tables** **9**

**3 Programs and Versions** **13**

**4 Other Sample and Laboratory Details** **14**

**8 Phylomorphospace Construction (Figure 2)** **29**

**9 Raw Sequence Data Availability** **30**

### 1 Materials and Methods

#### 1.1 Taxon sampling and sequencing

To generate sequencing data from specimens that were not preserved in ideal conditions and/or are more than five years post-collection (including a large library of previously sequenced chiton specimens from the personal collections of D.J. Eernisse as part of ongoing systematic revisions), we generated a suite of target capture loci. We designed target capture baits to predicted exon regions following a similar approach to Hugall et al. (52). We identified a set of 891 orthologs from 19 chiton transcriptomes and three reference molluscan genomes (Octopus bimaculoides, Lottia gigantea, and Crassostrea gigas) using the phylogenomics pipeline from Kocot et al. (53). We identified conserved exons from the three mollusc genomes using EnsemblMetazoa and predicted intron-exon boundaries in the chiton sequences with a custom script using tBLASTx. From this, we compiled a set of predicted chiton exons from 355 genes that had at least four representative chiton sequences from our transcriptome dataset. Arbor Biosciences synthesized 19,980 biotinylated DNA baits that target these regions. The baits have a length of 120 bp and a GC content between 35-55

We used the MYbaits Custom DNA-Seq kit v. 3.02 (Arbor Biosciences, Ann Arbor, Michigan, USA) according to manufacturer's instructions. We applied our target capture baits to DNA extracted from chiton specimens preserved in ethanol. We extracted total DNA from 87 specimens of preserved chitons with a phenol-chloroform extraction. We then produced Illumina sequencing libraries with unique barcodes using the NEBNext® Ultra™ II FS Library Prep Kit (New England Biolabs, Ipswich, Massachusetts, USA) with a 25 minute incubation at the fragmentation step. We used the TapeStation (Agilent, Santa Clara, California, USA) automated electrophoresis to confirm that the libraries had an average fragment size of 250 bp with adapters and that adapter dimers were absent. We incubated these libraries with biotinylated DNA baits via the MYbaits Custom DNA-Seq kit, and we pulled down the baits that hybridized with the Illumina library fragments with a streptavidin-coated magnetic bead binding process before washing away the uncaptured library fragments. We amplified the captured fragments with the KAPA HiFi Library Amp Kit (Roche Molecular Systems, Inc., Wilmington, Massachusetts, USA.) After the amplification, we confirmed that the libraries maintained an average fragment size of 250 bp and that adapter dimers were absent. We pooled enriched sequencing libraries and sequenced them on either an Illumina MiSeq (11 species) or NextSeq (68 species).

We trimmed Illumina adapters and low-quality sequences (quality score of ¡20) using TrimGalore v0.4.1 (54). We used HybPiper (55) to map reads to reference sequences. To expand our taxon sampling further, we combined target capture data with chiton transcriptomes from recent phylogenies of chitons that included chitons with only aesthete, chitons with eyespots, and chitons with shell eyes (56). We also included transcriptomes of several aplacophoran molluscs and a few conchiferan molluscs to root the phylogeny (Accession numbers in Supplementary Material, Outgroup and Available Sequences). To recover our target capture loci from chiton transcriptomes, we aligned the sequencing results from each target capture locus across chitons and determined the consensus DNA sequence for each. Then, we created a local blast database of each new chiton transcriptome and used each consensus DNA sequence from a target-capture locus as a query to locate the corresponding sequence within the new transcriptome. For each gene that we included, we retrieved only a single hit at E-value threshold 1e-4, so we are confident in matching gene sequences.

| **Order** | **Suborder** | **Family** | **Genus** | **Species** | **SampleID** | **Museum** | **Authority** | **Accession** | **Collection localality** |
|---|---|---|---|---|---|---|---|---|---|
| Order | Suborder | Family* | Genus* | Species* | Authority | SampleID | Museum** | Accession | Collection locality |
| Lepidopleurida | Lepidopleurina | Hanleyidae | Hanleya | hanleyi | (Bean, 1844) | HNAG | | SRX8235059 | Norway: Bergen |
| Lepidopleurida | Lepidopleurina | Leptochitonidae | Leptochiton | asellus | (Gmelin, 1791) | LASE | | SRX8235060 | Northeastern Atlantic Ocean |
| Lepidopleurida | Lepidopleurina | Leptochitonidae | Leptochiton | cf. rugatus | Carpenter, 1864 | LRUG | | SRX731464 | WA, USA |
| Lepidopleurida | Lepidopleurina | Leptochitonidae | Leptochiton | nexus | (Carpenter, 1892) | de8722 | xxxx | | San Pedro, California, USA |
| Lepidopleurida | Lepidopleurina | Nierstrazellidae | Nierstraszella | cf. lineata | (Nierstrasz, 1905) | de3358 | CASIZ-187921 | | Luzon Island, Philippines |
| Lepidopleurida | Lepidopleurina | Protochitonidae | Oldroydia | percrassa | (Dall, 1894) | de7537 | xxxx | | San Pedro, CA, USA |
| Callochitonida | | Callochitonidae | Callochiton | bouveti | Thiele, 1906 | de0106 | xxxx | | Bouvetoya Id., S. Atlantic |
| Callochitonida | | Callochitonidae | Callochiton | crocinus | (Reeve, 1847) | de8409 | MNHN-IM-2013-66754 | | Tasmania, Australia |
| Callochitonida | | Callochitonidae | Callochiton | dentatus | (Spengler, 1797) | de7501 | xxxx | | Chintsa West, Eastern Cape, South Africa |
| Callochitonida | | Callochitonidae | Callochiton | sp. | | CALL | | SRX8235057 | Ross Sea, Antarctica |
| Chitonida | Acanthochitonina | Acanthochitonidae | Acanthochitona | fascicularis | (Linnaeus, 1767) | ACRI | | SRX2422912 | Roscoff, France |
| Chitonida | Acanthochitonina | Acanthochitonidae | Cryptoconchus | floridanus | (Dall, 1889) | de8407 | MNHN-IM-2013-66590 | | Guadeloupe, Lesser Antilles Is. |
| Chitonida | Acanthochitonina | cf. Acanthochitonidae | Nuttallochiton | mirandus | (Thiele, 1906) | NMIR | | SRX8235049 | Weddell Sea, Antarctica |
| Chitonida | Acanthochitonina | Lepidochitonidae | Cyanoplax | dentiens | (Gould, 1846) | de8010 | xxxx | | Monterey Co., CA, USA |
| Chitonida | Acanthochitonina | Mopaliidae | Cryptochiton | stelleri | (Middendorff, 1847) | Cstelleri | | DRX136137 | Monterey, CA, USA |
| Chitonida | Acanthochitonina | Mopaliidae | Katharina | tunicata | (Wood, 1815) | Ktunicata | | SRX8235051 | Cattle Point, Friday Harbor, WA, USA |
| Chitonida | Acanthochitonina | Mopaliidae | Mopalia | hindsii | (Reeve, 1847) | de8020 | xxxx | | San Luis Obispo Co., CA, USA |
| Chitonida | Acanthochitonina | Mopaliidae | Mopalia | muscosa | (Gould, 1846) | MMUS | | | Santa Barbara, California, USA |
| Chitonida | Acanthochitonina | Mopaliidae | Mopalia | muscosa | (Gould, 1846) | Mmuscosa | | SRX8144968 | Cattle Point, Friday Harbor, WA, USA |
| Chitonida | Acanthochitonina | Mopaliidae | Tonicella | lineata | (Wood, 1815) | TLIN | | SRX8145069 | Cattle Point, Friday Harbor, WA, USA |
| Chitonida | Acanthochitonina | Plaxiphoridae | Plaxiphora | albida | (Blainville, 1825) | PALB | | | near Freemantle, WA, Australia |
| Chitonida | Acanthochitonina | Plaxiphoridae | Plaxiphora | aurata | (Spalowsky, 1795) | de7357 | xxxx | | Punta Arenas, Chile |
| Chitonida | Acanthochitonina | Plaxiphoridae | Plaxiphora | matthewsi | Iredale, 1910 | de8696 | xxxx | | San Remo, Victoria, Australia |
| Chitonida | Chitonina | Callistoplacidae | Callistochiton | ashbyi | (Barnard, 1963) | de2912 | xxxx | | Lavanono, Madagascar |
| Chitonida | Chitonina | Callistoplacidae | Callistochiton | cf. antiquus | (Reeve, 1847) | de8695 | xxxx | | San Remo, Victoria, Australia |
| Chitonida | Chitonina | Callistoplacidae | Callistochiton | palmulatus | Carpenter [in Dall], 1879 | de8013 | xxxx | | Monterey Co., CA, USA |
| Chitonida | Chitonina | Callistoplacidae | Callistochiton | shuttleworthianus | Pilsbry, 1893 | de8005 | xxxx | | Akumal, Q.R., Mexico |
| Chitonida | Chitonina | Callistoplacidae | Callistochiton | sp. A | | de8717 | xxxx | | La Paz, B.C.S., Mexico |
| Chitonida | Chitonina | Callistoplacidae | Callistochiton | sp. B | | de7491 | xxxx | | Bahía de Los Angeles, B.C., Mexico |
| Chitonida | Chitonina | Callistoplacidae | Calloplax | janeirensis | (Gray, 1828) | de8408 | MNHN-IM-2013-66596 | | Guadeloupe, Lesser Antilles Is. |
| Chitonida | Chitonina | Callistoplacidae | Calloplax | vivipara | (Plate, 1899) | de6100 | xxxx | | El Tabo, Valparaíso, Chile |
| Chitonida | Chitonina | Chaetopleuridae | Chaetopleura | apiculata | (Say, 1834) | CAPI | | SRX8235058 | Wood's Hole, MA, USA |
| Chitonida | Chitonina | Chaetopleuridae | Chaetopleura | benaventei | Plate, 1899 | de7444 | xxxx | | Hualpén, near Concepción, Chile |
| Chitonida | Chitonina | Chaetopleuridae | Chaetopleura | lurida | (G. B. Sowerby I, 1832) | de8719 | xxxx | | La Paz, B.C.S., Mexico |
| Chitonida | Chitonina | Chaetopleuridae | Chaetopleura | peruviana | (Lamarck, 1819) | de6126 | xxxx | | Huasco, Atacama, Chile |
| Chitonida | Chitonina | Chitonidae | Acanthocpleura | echinata | (Barnes, 1824) | de6124 | xxxx | | Lobitos, Perú |
| Chitonida | Chitonina | Chitonidae | Acanthopleura | gemmata | (Blainville, 1825) | de3230 | xxxx | | Ningaloo, WA, Australia |
| Chitonida | Chitonina | Chitonidae | Acanthopleura | gemmata | (Blainville, 1825) | de6119 | xxxx | | Seragaki, Okinawa Id., Japan |
| Chitonida | Chitonina | Chitonidae | Acanthopleura | granulata | (Gmelin, 1791) | de3493 | | | Keys, FL, USA |
| Chitonida | Chitonina | Chitonidae | Acanthopleura | planispina | Bergenhayn, 1933 | de2429 | xxxx | | Ogasawara Is., Japan |
| Chitonida | Chitonina | Chitonidae | Acanthopleura | spinosa | (Bruguière, 1792) | de3229 | xxxx | | Ningaloo, WA, Australia |
| Chitonida | Chitonina | Chitonidae | Chiton | cumingsii | Frembly, 1827 | de6125 | xxxx | | Huasco, Atacama, Chile |
| Chitonida | Chitonina | Chitonidae | Chiton | marmoratus | Gmelin, 1791 | CMAR | | SRX8235047 | St. Thomas Id., U.S. Virgin Is. |
| Chitonida | Chitonina | Chitonidae | Chiton | tuberculatus | Linnaeus, 1758 | CTUB | | SRX8235048 | St. Thomas Id., U.S. Virgin Is. |
| Chitonida | Chitonina | Chitonidae | Chiton | virgulatus | G. B. Sowerby II, 1840 | de7506 | xxxx | | La Paz, B.C.S., Mexico |
| Chitonida | Chitonina | Chitonidae | Chiton | viridis | Spengler, 1797 | de6033 | xxxx | | Akumal, Q.R., Mexico |
| Chitonida | Chitonina | Chitonidae | Enoplochiton | niger | (Barnes, 1824) | de6128 | xxxx | | Huasco, Atacama, Chile |
| Chitonida | Chitonina | Chitonidae | Liolophura | japonica | (Lischke, 1873) | de8602 | xxxx | | Jeju Id., S. Korea |
| Chitonida | Chitonina | Chitonidae | Lucilina | lamellosa | (Quoy and Gaimard, 1835) | de8404 | MNHN-IM-2013-14546 | | Papua New Guinea |
| Chitonida | Chitonina | Chitonidae | Onithochiton | neglectus | Rochebrune, 1881 | de7226 | xxxx | | Wellington, New Zealand |
| Chitonida | Chitonina | Chitonidae | Onithochiton | noemiae | (Rochebrune, 1884) | ONOE | | | Kanumera Bay,New Caledonia, France |
| Chitonida | Chitonina | Chitonidae | Onithochiton | quercinus | (Gould, 1846) | de3369 | xxxx | | Point Cartwright, QLD, Australia |
| Chitonida | Chitonina | Chitonidae | Onithochiton | sp. A | | de8402 | xxxx | | near Jakarta, Indonesia |
| Chitonida | Chitonina | Chitonidae | Onithochiton | sp. B | | de8006 | UF457518 | | Saipan Island; Northern Mariana Islands, Guam, USA |
| Chitonida | Chitonina | Chitonidae | Radsia | barnesii | Gray, 1828 | de7502 | xxxx | | Caldera, Atacama, Chile |
| Chitonida | Chitonina | Chitonidae | Rhyssoplax | calliozona | (Pilsbry, 1894) | de6123 | xxxx | | near Adelaide, SA, Australia |
| Chitonida | Chitonina | Chitonidae | Rhyssoplax | maldivensis | (E. A. Smith, 1903) | de8018 | UF463612 | | Farasan Banks, Saudi Arabia |
| Chitonida | Chitonina | Chitonidae | Rhyssoplax | olivacea | (Spengler, 1797) | ROLI | | SRX205322 | |
| Chitonida | Chitonina | Chitonidae | Squamopleura | araucariana | (Hedley, 1898) | SARA | | | Kanumera Bay,New Caledonia, France |
| Chitonida | Chitonina | Chitonidae | Squamopleura | cf. miles | (Pilsbry, 1894) | de8401 | xxxx | | near Jakarta, Indonesia |
| Chitonida | Chitonina | Chitonidae | Sypharochiton | pelliserpentis | (Quoy and Gaimard, 1835) | de7232 | xxxx | | Wellington, New Zealand |
| Chitonida | Chitonina | Chitonidae | Tegulaplax | hululensis | (E. A. Smith, 1903) | de3386 | xxxx | | Kyan, Okinawa Id., Japan |
| Chitonida | Chitonina | Chitonidae | Tonicia | calbucensis | Plate, 1897 | de6127 | xxxx | | Huasco, Atacama, Chile |
| Chitonida | Chitonina | Chitonidae | Tonicia | forbesii | Carpenter, 1857 | | xxxx | | King George Id., Shetland Is. |
| Chitonida | Chitonina | Chitonidae | Tonicia | lebruni | Rochebrune, 1884 | de7353 | xxxx | | Punta Arenas, Chile |
| Chitonida | Chitonina | Chitonidae | Tonicia | schrammi | (Shuttleworth, 1856) | TONI | | SRX8235050 | |
| Chitonida | Chitonina | Ischnochitonidae | Ischnochiton | bouryi | Dupuis, 1917 | de8405 | MNHN-IM-2013-16841 | | Papua New Guinea |
| Chitonida | Chitonina | Ischnochitonidae | Ischnochiton | comptus | (Gould, 1859) | de8605 | xxxx | | Jeju Id., S. Korea |
| Chitonida | Chitonina | Ischnochitonidae | Ischnochiton | erythronotus | (C. B. Adams, 1845) | de6985 | xxxx | | St. Thomas Id., U.S. Virgin Is. |
| Chitonida | Chitonina | Ischnochitonidae | Ischnochiton | hakodadensis | Carpenter, 1893 | de3565 | xxxx | | Vostok Bay, Russia |
| Chitonida | Chitonina | Ischnochitonidae | Ischnochiton | lentiginosus | (G. B. Sowerby II, 1840) | de8285 | xxxx | | Mallacoota, Victoria, Australia |
| Chitonida | Chitonina | Ischnochitonidae | Ischnochiton | maorianus | Iredale, 1914 | de7228 | xxxx | | Wellington, New Zealand |

| | | | | | | | | |
|---|---|---|---|---|---|---|---|---|
| Chitonida | Chitonina | Ischnochitonidae | Ischnochiton | oniscus | (Krauss, 1848) | de8007 | xxxx | Port Alfred, Eastern Cape, South Africa |
| Chitonida | Chitonina | Ischnochitonidae | Ischnochiton | pusio | (G. B. Sowerby I, 1832) | de8014 | xxxx | Huasco, Atacama, Chile |
| Chitonida | Chitonina | Ischnochitonidae | Ischnochiton | sirenkoi | Dell'Angelo et al., 2011 | de2914 | xxxx | Lavanono, Madagascar |
| Chitonida | Chitonina | Ischnochitonidae | Ischnochiton | sp. | | de8016 | xxxx | Nahoon Pt., E. London, South Africa |
| Chitonida | Chitonina | Ischnochitonidae | Ischnochiton | stramineus | (G. B. Sowerby I, 1832) | de7359 | xxxx | Punta Arenas, Chile |
| Chitonida | Chitonina | Ischnochitonidae | Ischnochiton | tridentatus | Pilsbry, 1893 | de8015 | xxxx | La Paz, B.C.S., Mexico |
| Chitonida | Chitonina | Ischnochitonidae | Ischnochiton | versicolor | (G. B. Sowerby II, 1840) | de8410 | MNHN-IM-2013-66756 | Tasmania, Australia |
| Chitonida | Chitonina | Ischnochitonidae | Ischnochiton | winckworthi | Leloup, 1936 | de2440 | xxxx | Phi Phi Don, Krabi Province, Thailand |
| Chitonida | Chitonina | Ischnochitonidae | Ischnochiton | yerburyi | (E. A. Smith, 1891) | de8406 | MNHN-IM-2013-66552 | Inhaca Id., Mozambique Channel, Mozambique |
| Chitonida | Chitonina | Ischnochitonidae | Ischnoplax | petctinata | (G. B. Sowerby II, 1840) | de8716 | MNHN-IM-2013-66192 | Guadeloupe, Lesser Antilles Is. |
| Chitonida | Chitonina | Ischnochitonidae | Lepidozona | clathrata | (Reeve, 1847) | de7507 | xxxx | La Paz, B.C.S., Mexico |
| Chitonida | Chitonina | Ischnochitonidae | Lepidozona | mertensii | (Middendorff, 1847) | de8720 | UF476217 | San Juan Island., WA, USA |
| Chitonida | Chitonina | Ischnochitonidae | Lepidozona | radians | (Carpenter, 1892) | de7499 | xxxx | Monterey Co., CA, USA |
| Chitonida | Chitonina | Ischnochitonidae | Lepidozona | retiporosa | (Carpenter, 1864) | de8019 | xxxx | San Pedro, CA, USA |
| Chitonida | Chitonina | Ischnochitonidae | Lepidozona | serrata | (Carpenter, 1864) | de8017 | xxxx | La Paz, B.C.S., Mexico |
| Chitonida | Chitonina | Ischnochitonidae | Lepidozona | willetti | (Berry, 1917) | de8721 | xxxx | Monterey Co., CA, USA |
| Chitonida | Chitonina | Ischnochitonidae | Stenochiton | cymodocealis | Ashby, 1918 | de8012 | xxxx | Noble Rocks, Victoria, Australia |
| Chitonida | Chitonina | Ischnochitonidae | Stenoplax | conspicua | (Dall, 1879) | SCON | | San Onofre, California, USA |
| Chitonida | Chitonina | Ischnochitonidae | Stenoplax | limaciformis | (G. B. Sowerby I, 1832) | de2781 | xxxx | La Paz, B.C.S., Mexico |
| Chitonida | Chitonina | Ischnochitonidae | Stenoplax | mariposa | (Dall, 1919) | de3461 | xxxx | La Paz, B.C.S., Mexico |
| Chitonida | Chitonina | Ischnochitonidae | Stenosemus | mexicanus | (Kaas, 1993) | de4903 | xxxx | Gulf of Mexico |
| Chitonida | Chitonina | Ischnochitonidae | Tonicina | zschaui | Carpenter, 1857 | de7505 | xxxx | Mazatlan, Sinoloa, Mexico |
| Chitonida | Chitonina | Ischnochitonidae | Tonicina | zschaui | (Pfeffer, 1886) | de8093 | xxxx | King George Id., Shetland Is. |
| Chitonida | Chitonina | Ischnochitonidae | Tripoplax | trifida | (Carpenter, 1864) | de8460 | xxxx | San Juan Island, WA, USA |
| Chitonida | Chitonina | Loricidae | Lorica | volvox | (Reeve, 1847) | de8088 | xxxx | Tasmania, Australia |
| Chitonida | Chitonina | Loricidae | Loricella | angasi | (H. Adams, 1864) | de8294 | xxxx | Stony Point, Victoria, Australia |
| Chitonida | Chitonina | Schizochitonidae | Schizochiton | incisus | (G. B. Sowerby II, 1841) | de8403 | MNHN-IM-2013-14442 | Papua New Guinea |

*Most taxa follow MolluscaBase; all differences are intentional and reflect ongoing systematic revisions by DJE
**Note to reviewers: The 'xxxx' placeholders in this column will be updated in a subsequent version of this manuscript when museum catalog numbers become available.

#### 1.2 Phylogeny

To generate a robust phylogeny of chitons, we compared coalescence and concatenation-based approaches using Maximum Likelihood, and Bayesian approaches. We first aligned all loci with MAFFT v7.305 (58) to produce a single data matrix of 103 chitons that could be partitioned by locus. We produced a maximum likelihood tree in IQ-TREE2 (57), partitioning by gene and using ModelFinder to determine the best model for each locus. We also calculated gene concordance factors in IQ-TREE2. Separately, we produced a maximum-likelihood phylogeny for each gene individually with IQ-TREE2, again using ModelFinder, and input these trees into ASTRAL-Pro (41), which estimates a species tree from multiple gene trees via the multispecies coalescence (MSC) model and a quartet-based approach.

The placement of *Schizochiton* has been uncertain across studies of chitons, and our study is limited by the inclusion of only a single representative of the genus *Schizochiton*, *Schizochiton incisus*. Specimens of the only other accepted congeneric species, *S. jousseaumei* (Dupuis, 1917) were not available, and this is the only genus in the superfamily, Schizochitonoidea.Some morphological characters, most notably pectinated insertion plates and adanal gills, suggest an affinity of *Schizochiton* with Chitonidae. However, differences of opinion persist as to whether these traits are useful apomorphies. For example, the insertion plates of *Schizochiton* were once termed "obsoletely pectinated" (59) although they have strong pectination comparable to members of Chitonidae (DJE, pers. observation). Additionally, the caudal sinus is absent in Chitonidae but present in *Schizochiton* (65). However, caudal sinuses evolved multiple times across chitons, so the presence or absence of a sinus may not be a useful morphological character. *Schizochiton* has been recovered as sister to the remaining Chitonina, but with weak support (36). Our placement of *Schizochiton* requires that pectinated insertion plates evolved convergently in *Schizochiton* and Chitonidae. To further investigate the placement of *Schizochiton* in our phylogeny, we used an iterative pruning method to verify that *Schizochiton* was not an unstable branch (90) and determined that *Schizochiton* did not have any indicators of being a long branch or a highly unstable branch (Supplemental Data, RogueNaRok).

#### 1.3 Ancestral state reconstruction and morphospace determination

To determine how many times distributed visual systems evolved across the chiton phylogeny, we performed ancestral state reconstruction using RayDISC and corHMM in R, which take a tree file and a corresponding character matrix for a given trait and estimates transition rates and ancestral states for binary or multistate traits (Beaulieu and O'Meara 2014). We used the consensus tree file from our MrBayes dated phylogeny run with 15 loci from sortadate as our input tree, and inferred ancestral states for shell eyes, eyespots, and the maximum number of insertion slits on the anteriormost shell plate. We gathered information on slit number from volumes of Kaas's monographs (60-64). We took the highest recorded slit number for the anterior valve to account for the impact of animal size on slit number, where slit number can increase across growth.

| Species | Slits | Citation | Notes |
|---|---|---|---|
| Acanthochitona fascicularis | 5 | Kaas 1985 | |
| Acanthopleura gemmata | 11 | Kaas, Van Belle, and Strack 2006 | |
| Acanthopleura gemmata2 | 11 | Kaas, Van Belle, and Strack 2006 | |
| Acanthopleura granulata | 12 | Brooker 2003 | |
| Acanthopleura planispina | 10 | Kaas, Van Belle, and Strack 2006 | |
| Acanthopleura spinosa | 12 | Brooker 2003 | |
| Apodomenia enigmatica | | | |
| Callistochiton antiquus B | 10 | Kaas and Van Belle 1994b | |
| Callistochiton ashbyi | 9 | Barnard K.H. 1963 | |
| Callistochiton elenensis A | 9 | Kaas and Van Belle 1994b | |
| Callistochiton palmulatus | 11 | Kaas and Van Belle 1994b | |
| Callistochiton shuttleworthianus | 11 | Kaas and Van Belle 1994b | |
| Callochiton bouveti | 16 | Kaas and Van Belle 1985b | |
| Callochiton crocinus | 20 | Kaas and Van Belle 1985b | |
| Callochiton dentatus | 25 | Kaas and Van Belle 1985b | |
| Callochiton sp. | | | |
| Calloplax janeirensis | 10 | Kaas and Van Belle 1994b | |
| Calloplax vivipara | 9 | Kaas and Van Belle 1994b | |

| | | | |
|---|---|---|---|
| Chaetopleura apiculata | 12 | Kaas, Van Belle, and Strack 2006 | |
| Chaetopleura benaventei | 11 | Kaas and Van Belle 1987 | |
| Chaetopleura lurida | 11 | Kaas and Van Belle 1987 | |
| Chaetopleura peruviana | 10 | Kaas and Van Belle 1987 | |
| Chiton cumingsii | 14 | Kaas, Van Belle, and Strack 2006 | |
| Chiton marmoratus | 15 | Kaas, Van Belle, and Strack 2006 | |
| Chiton tuberculatus | 15 | Kaas, Van Belle, and Strack 2006 | |
| Chiton virgulatus | 20 | Kaas, Van Belle, and Strack 2006 | |
| Chiton viridis | 14 | Kaas, Van Belle, and Strack 2006 | |
| Cryptoconchus floridanus | 5 | Reyes-Gomez 2017 | |
| Chrysomallon squamiferum | | | |
| Crassostrea virginica | | | |
| Cryptochiton stelleri | 7 | Sirenko 2006 | |
| Cyanoplax dentiens | 11 | Kaas and Van Belle 1985b | |
| Callistochiton sp.B | | | |
| Acanthopleura echinata | 8 | Kaas, Van Belle, and Strack 2006 | |
| Enoplochiton niger | 9 | Kaas, Van Belle, and Strack 2006 | |
| Hanleya hanleyi | 0 | Kaas and Van Belle 1985a | |
| Ischnochiton bouryi | 11 | Kaas and Van Belle 1994a | |
| Ischnochiton comptus | 13 | Gould, A. A. 1859 | |
| Ischnochiton erythronotus | 10 | Kaas and Van Belle 1994a | |
| Ischnochiton hakodadensis | 20 | Kaas and Van Belle 1994a | |
| Ischnochiton lentiginosus | 12 | Kaas and Van Belle 1994b | |
| Ischnochiton maorianus | 12 | Kaas and Van Belle 1994a | |
| Ischnochiton oniscus | 13 | Kaas and Van Belle 1994a | |
| Ischnochiton pusio | 14 | Kaas and Van Belle 1994b | |
| Ischnochiton sp. | | | |
| Ischnochiton sirenkoi | 11 | Dell'Angelo et al. 2011 | |
| Ischnochiton stramineus | 15 | Kaas and Van Belle 1994a | |
| Ischnochiton tridentatus | 18 | Kaas and Van Belle 1994a | |
| Ischnochiton versicolor | 13 | Kaas and Van Belle 1994a | |
| Ischnochiton winckworthi | 11 | Kaas and Van Belle 1994a | |
| Ischnochiton yerburyi | 11 | Kaas and Van Belle 1994a | |
| Ischnoplax sp. | | | |
| Katharina tunicata | 8 | Kaas and Van Belle 1994b | |
| Lepidozona clathrata | 11 | Kaas and Van Belle 1987 | mode used |
| Lepidozona mertensii | 12 | Kaas and Van Belle 1987 | |
| Lepidozona radians | 11 | Pilsbry, H. A. 1892-1893 | |
| Lepidozona retiporosa | 11 | Kaas and Van Belle 1987 | mode used |
| Lepidozona serrata | 12 | Kaas and Van Belle 1987 | |
| Lepidozona willetti | 11 | Kaas and Van Belle 1987 | |
| Leptochiton asellus | 0 | Kaas and Van Belle 1985a | |
| Leptochiton nexus | 0 | Kaas and Van Belle 1985a | |
| Leptochiton rugatus | 0 | Kaas and Van Belle 1985a | |
| Liolophura japonica | 11 | Kaas, Van Belle, and Strack 2006 | |
| Lorica volvox | 9 | Kaas, Van Belle, and Strack 2006 | |
| Loricella angasi | 8 | Kaas, Van Belle, and Strack 2006 | |
| Lucilina lamellosa | 8 | Kaas, Van Belle, and Strack 2006 | |
| Mopalia muscosa2 | 9 | Kaas and Van Belle 1994b | |
| Mopalia hindsii | 8 | Kaas and Van Belle 1994b | |
| Mopalia muscosa | 9 | Kaas and Van Belle 1994b | |
| Onithochiton spGuam | | | |
| Nierstraszella cf.lineata | 0 | Kaas and Van Belle 1985a | |
| Nuttallochiton mirandus | 10 | Kaas and Van Belle 1987 | |
| Oldroydia percrassa | 0 | Kaas and Van Belle 1985a | |
| Onithochiton sp.B Indonesia | | | |
| Onithochiton neglectus | 8 | Kaas, Van Belle, and Strack 2006 | |
| Onithochiton noemiae | 8 | Kaas, Van Belle, and Strack 2006 | |

| | | |
|---|---|---|
| Onithochiton quercinus | 8 | Kaas, Van Belle, and Strack 2006 |
| Plaxiphora albida | 8 | Kaas and Van Belle 1994b |
| Plaxiphora aurata | 8 | Kaas and Van Belle 1994b |
| Plaxiphora matthewsi | 8 | Kaas and Van Belle 1994b |
| Proneomenia sluiteri | | |
| Radsia barnesii | 16 | Kaas, Van Belle, and Strack 2006 |
| Rhyssoplax calliozona | 8 | Kaas, Van Belle, and Strack 2006 |
| Rhyssoplax maldivensis | 10 | Kaas, Van Belle, and Strack 2006 |
| Rhyssoplax olivacea | 9 | Kaas, Van Belle, and Strack 2006 |
| Schizochiton incisus | 7 | Kaas, Van Belle, and Strack 2006 |
| Squamopleura araucariana | 9 | Kaas, Van Belle, and Strack 2006 |
| Squamopleura sp. Indonesia | | |
| Stenochiton cymodocealis | 13 | Kaas and Van Belle 1994b |
| Stenoplax conspicua | 12 | Kaas and Van Belle 1987 |
| Stenoplax limaciformis | 13 | Kaas and Van Belle 1987 |
| Stenoplax mariposa | 10 | Kaas and Van Belle 1994a |
| Stenosemus mexicanus | 13 | Kaas 1993 |
| Scutopus ventrolineatus | | |
| Sypharochiton pelliserpentis | 13 | Kaas, Van Belle, and Strack 2006 |
| Tegulaplax hululensis | 10 | Kaas, Van Belle, and Strack 2006 |
| Tonicella lineata | 10 | Kaas and Van Belle 1985b |
| Tonicia calbucensis | 9 | Kaas, Van Belle, and Strack 2006 |
| Tonicia forbesii | 9 | Kaas, Van Belle, and Strack 2006 |
| Tonicia lebruni | 9 | Kaas, Van Belle, and Strack 2006 |
| Tonicia schrammi | 10 | Kaas, Van Belle, and Strack 2006 |
| Tonicina zschaui | 14 | Kaas and Van Belle 1985b |
| Tripoplax trifida | 13 | Kaas and Van Belle 1987 |

To estimate the density of aesthetes on a given shell plate, we used Fiji (Schindelin et al. 2012) to simplify SEM images of chiton plates by applying filters for cell counting from various sources (67-80). We then manually took a series of 10 estimates of shell eye, eyespot, and/or aesthete density (aesthetes/mm2, including both micraesthetes + macraesthetes) per animal (Supplemental Data, Aesthete Quantifications).

#### 1.4 Molecular dating analyses:

Following guidelines by Wolfe et al. for fossil calibrations across deep phylogenies, we selected fossil calibration points that span the depth of the phylogeny and are supported by multiple dating methods (82-83). There is a known bias in the chiton fossil record because members of Chitonida have thicker plates than Lepidopleurida (84), so there are more available calibration points for shallower nodes in the tree, and recent fossils are more easily identified (48,37). We set a hard maximum date for the entire phylogeny, including the outgroup Crassostrea virginica, of 549 million years ago, based on the Ediacaran deposits of the Nama group, a marine community with many of the earliest marine animal remains (including calcareous algae), but no remains of molluscs (39, 48). We assigned a soft maximum date of 359 million years to Neoloricata as a clade, based on the earliest fossils with a recognizable articulamentum (the inner layer of a shell plate) and thus the earliest likely emergence of Neoloricata (73, 48). We used this same soft maximum for all calibration points within Neoloricata. We assigned the Lepidopleurida a uniform calibration of 359 to 201.3 mya based on fossils from the Jurassic (67,74). We selected 66 mya as a minimum for Chitonida in agreement with Irisarri 2014 and Puchalski 2008 that earlier fossils are ambiguous and not plentiful enough to constrain the divergence of Chitonida from other chitons, so we used a fossil from the Upper Cretaceous as the more recent but definitive calibration point (74, 37, 85, 48). We set the minimum for Acanthochitonina at 33.9 mya based on fossils of Plaxiphora (which we in turn calibrated at a minimum of 23.03 mya) and Mopalia (which we calibrated at 15 to 359 mya) according to assessments and comparisons of multiple fossils by Puchalski (48). We set the genus Tonicia at 33.9 mya based on the fossils that bracket the divergence of Toniciinae from Acanthopleurina (48). This suite of fossils spans the expected phylogeny without constraining any of our clades of interest for the evolution of chiton visual systems.

To estimate divergence times across chiton evolution, we used a Bayesian relaxed molecular clock in MrBayes (83). Bayesian analyses are time-intensive (83), so we selected 15 loci with sortadate (87), a program that selects a subset of loci for analysis based first on congruence with a phylogeny (in this case, our maximum likelihood phylogeny from ASTRAL-Pro), and second based on how well each locus adheres to a clock-like accumulation of changes across a phylogeny. To ensure that using a subset of genes did not change the resulting dates of the phylogeny in a way that changed our interpretation of visual system evolution in chitons, we re-ran a Bayesian dating tree using sets of 15, 25, and 30 genes from sortadate. Though chains did not converge in a reasonable time with 30 genes (standard deviation of split frequencies between two concurrent chains did not drop below 0.05), tree topologies and age estimates remained consistent. We used a fixed amino acid substitution model of GTR across our 15 (or 25/30) partitions, and we applied the IGR model with a preset prior of exp(10) and a preset clock rate prior of lognorm(-7, 0.6). We set the branch lengths prior based on the fossilization clock, and ran each analysis a minimum of one million generations, extending as needed to reach a standard deviation of split frequencies ¡0.05.

#### 2 Supplementary Figures and Tables

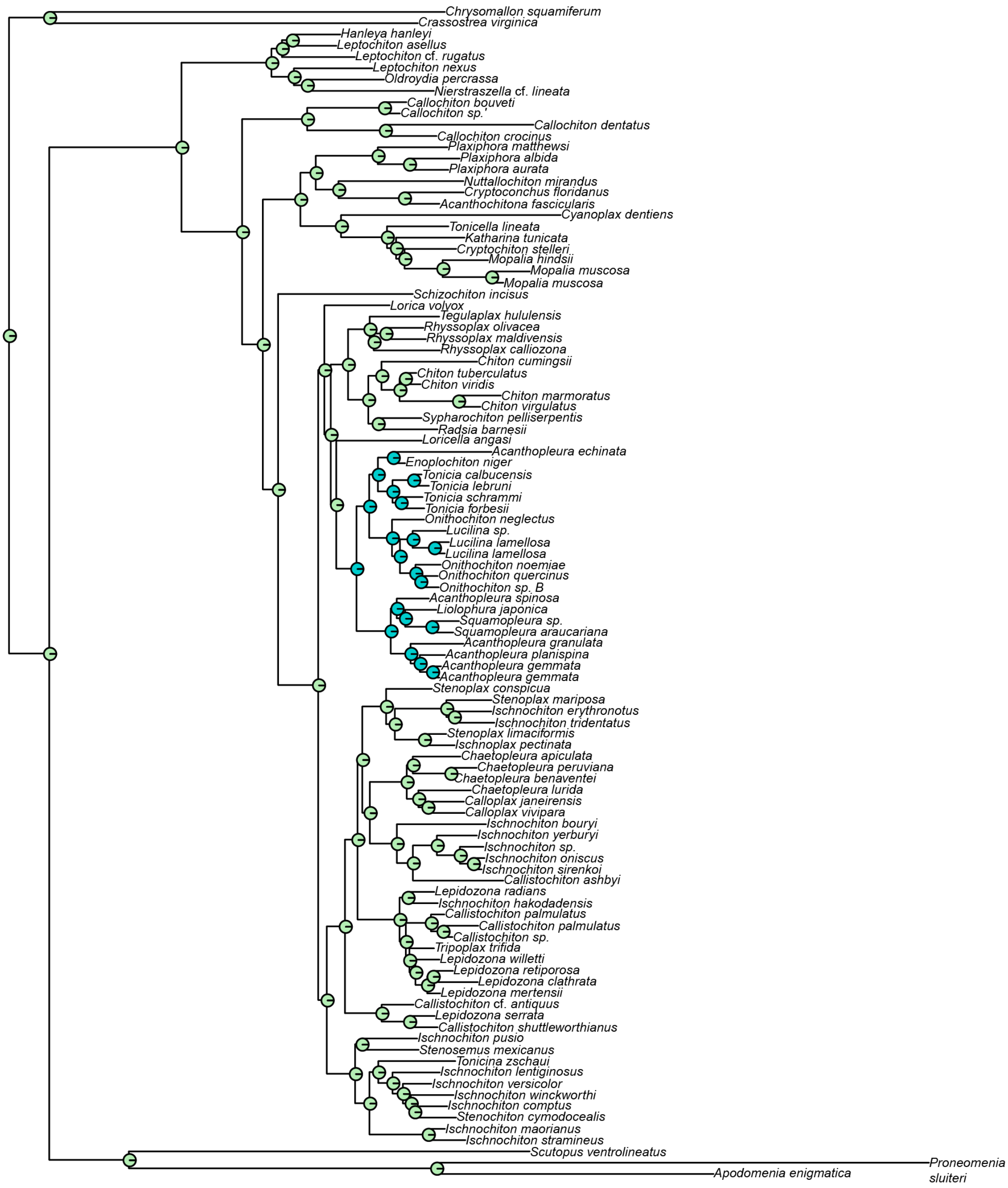

**Supplementary Fig. 1**: Ancestral state reconstruction of shell eyes shows ≥95% proportional marginal likelihood for two independent origins of shell eyes, one within the Chitonidae and one in /emphSchizochiton incisus (not visible because *S. incisus* is a monotypic branch). Blue circles indicate the presence of eyespots, green the absence of eyespots.

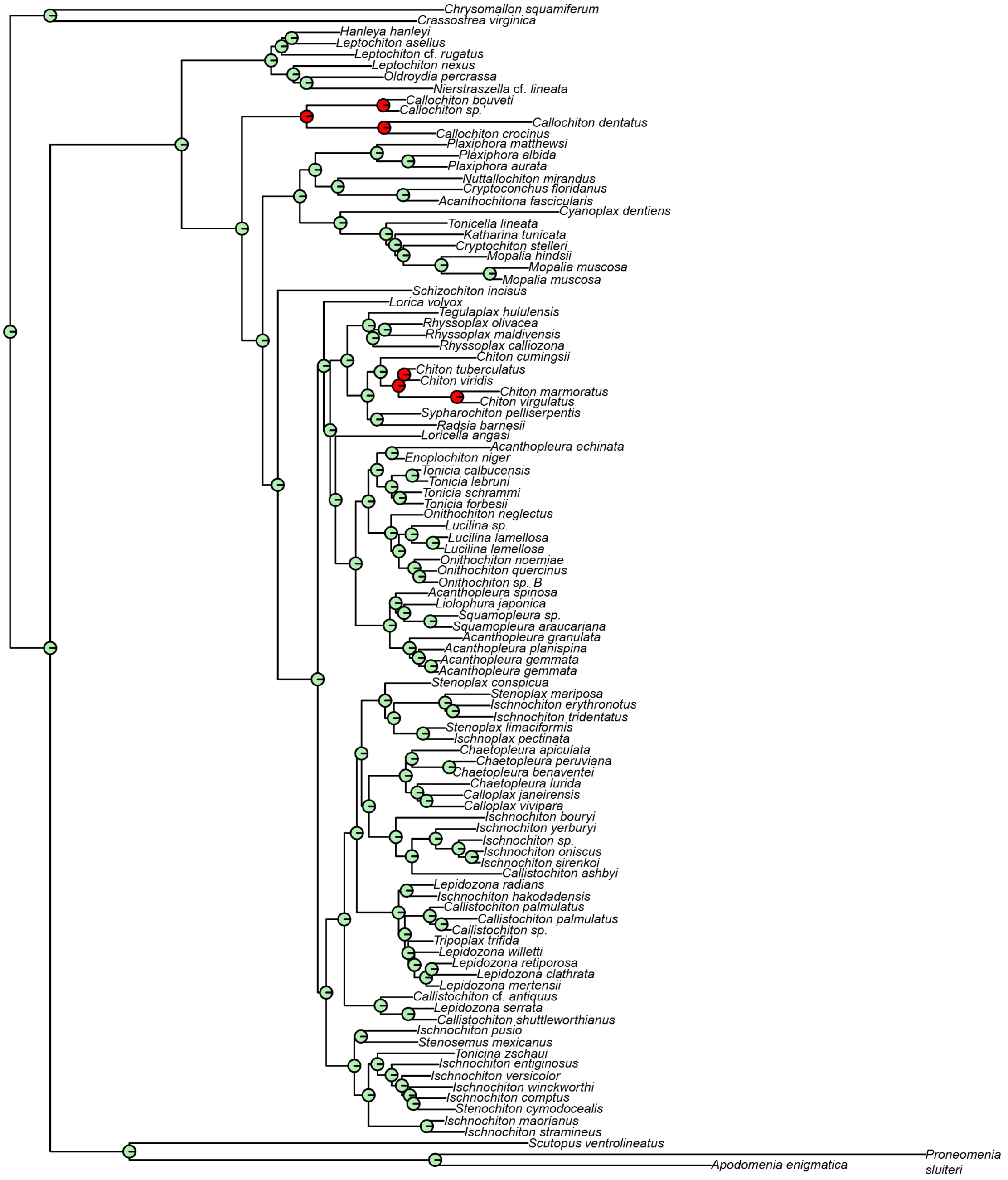

**Supplementary Fig. 2**: Ancestral state reconstruction of eyespots show ≥95% proportional marginal likelihood for two independent origins of eyespots, once in Callochitonida and once in Chitoninae. Red circles indicate the presence of eyespots, green the absence of eyespots.

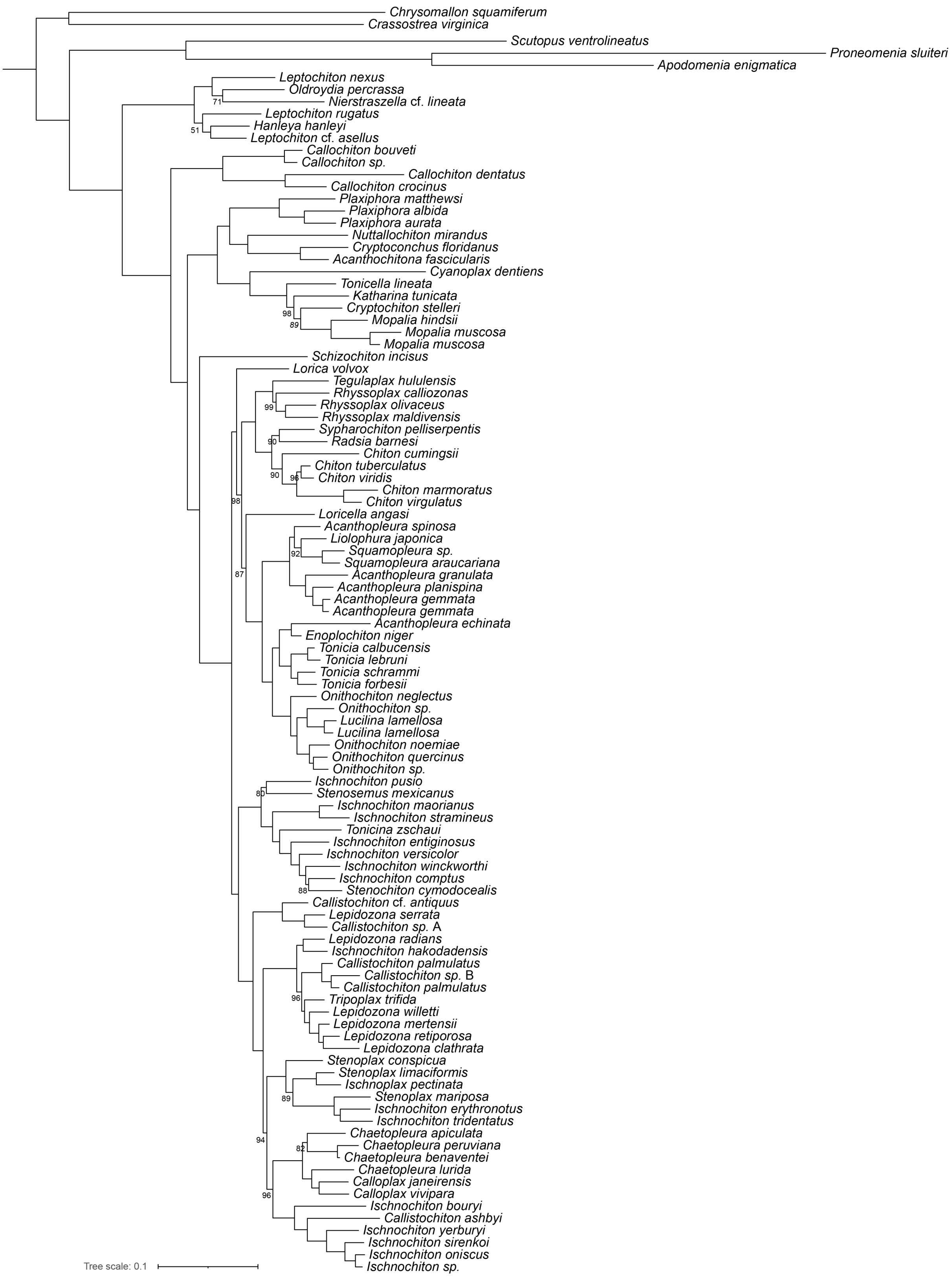

**Supplementary Fig. 3**: Maximum likelihood phylogeny of chitons based on the full matrix of all target capture loci generated in the present study. Tree made with IQTree2; all support values not shown are 100.

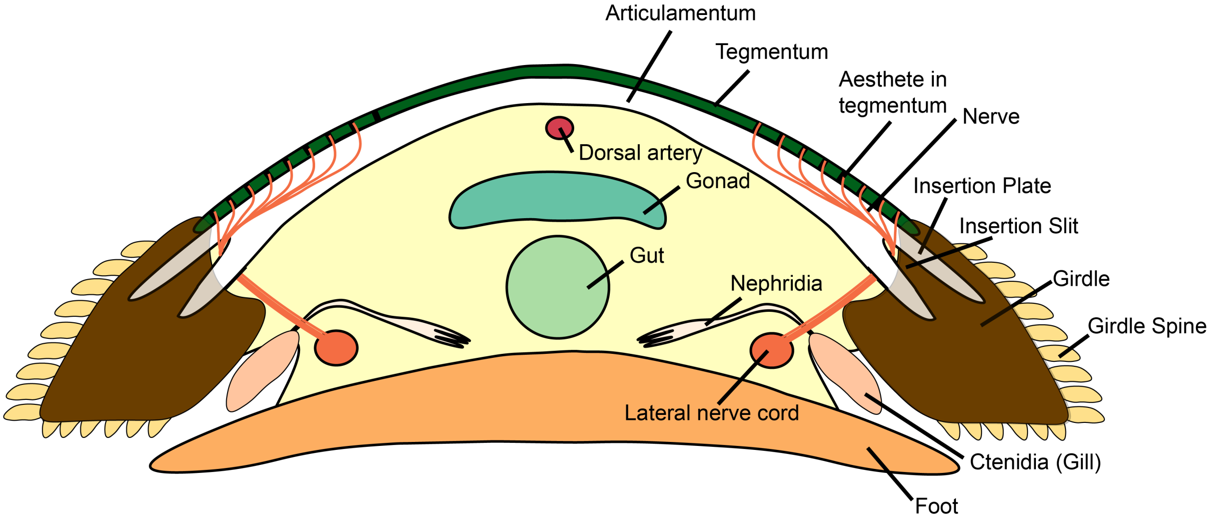

**Supplementary Fig. 4**: An axial cross section through a chiton with labeled parts, showing how nerves (orange) run through the tegmentum (green) and across the shell plate (white) to pass through an insertion slit between insertion plates to join the nerve cord below (orange). Insertion plates extend into the mantle (brown) to hold shell plates in place.

### 3 Programs and Versions

| Program | Version | Citation |
|---|---|---|
| IQ-TREE 2 | 2.0.3 | ((*16*)) |
| ASTRAL-PRO | 1.0 | ((*17*)) |
| RogueNaRok | Accessed May 2022 | ((*8*), (*11*)) |
| SortaDate | 1.0 | ((*15*)) |
| MrBayes | 3.2.6 | ((*9*)) |

### 4 Other Sample and Laboratory Details

The majority of samples available were plugs of foot tissue (muscle) preserved in ethanol. As such, we excluded from the analysis any samples for which the providence was not absolutely certain and/or the identification was unclear.

#### 4.1 Deconcatenating Initial Chiton Data

To pull out individual genes from additional transcriptomes, first partitioned data had to be de-concatenated using a custom script decat.py:

```
#decat.py Rebecca M Varney 2021
#First separating the names of the loci from the numeric range of each locus
loci=[]
names=[]
with open ("partitions.txt") as p:
    for line in p:
        line = line.strip ('LG, ')
        line = line.split (" = ")
        loci.append (line [1])
        names.append (line [0])

#Change the hyphens to colons
num_loci=[]
for string in loci:
    numloci = int(string.split("-")[0])-1 #Save the first number as an integer
    num_loci.append(numloci) # Add to list

# Open concatenated file
file = open("concatenated.fas")
c = "" # Empty string to save into
for line in file:
    c += line
file.close()
data = c.splitlines() # Save a data into a list broken up by lines

# Loop over loci names
for i in range(len(names)):
    # Create empty string to save data into for each loci
    sout = ""
    # Loop over data
    for line in data:
        if line[0] == ">":
            # Write name
            sout += line + "\n"
        else:
            if i == len(names)-1: # Need to do something special for the last loci to get the
                end of the string
                sout += line[num_loci[i]:] + "\n"
            else:
                sout += line[num_loci[i]:num_loci[i+1]] + "\n"
    #break
    ofile = open(names[i]+".fas",'w')
    ofile.write(sout)
    ofile.close()
```

#### 4.2 Aesthete Quantifications

To determine aesthete densities, we used available SEM images from as many species as we could find reliably. We used the cell counter function in FIJI (Image J) to determine the number of complete aesthetes (and eyespots and shell eyes) visible within a frame. Below is a midway example, with macraesthetes and micraestehtes counted:

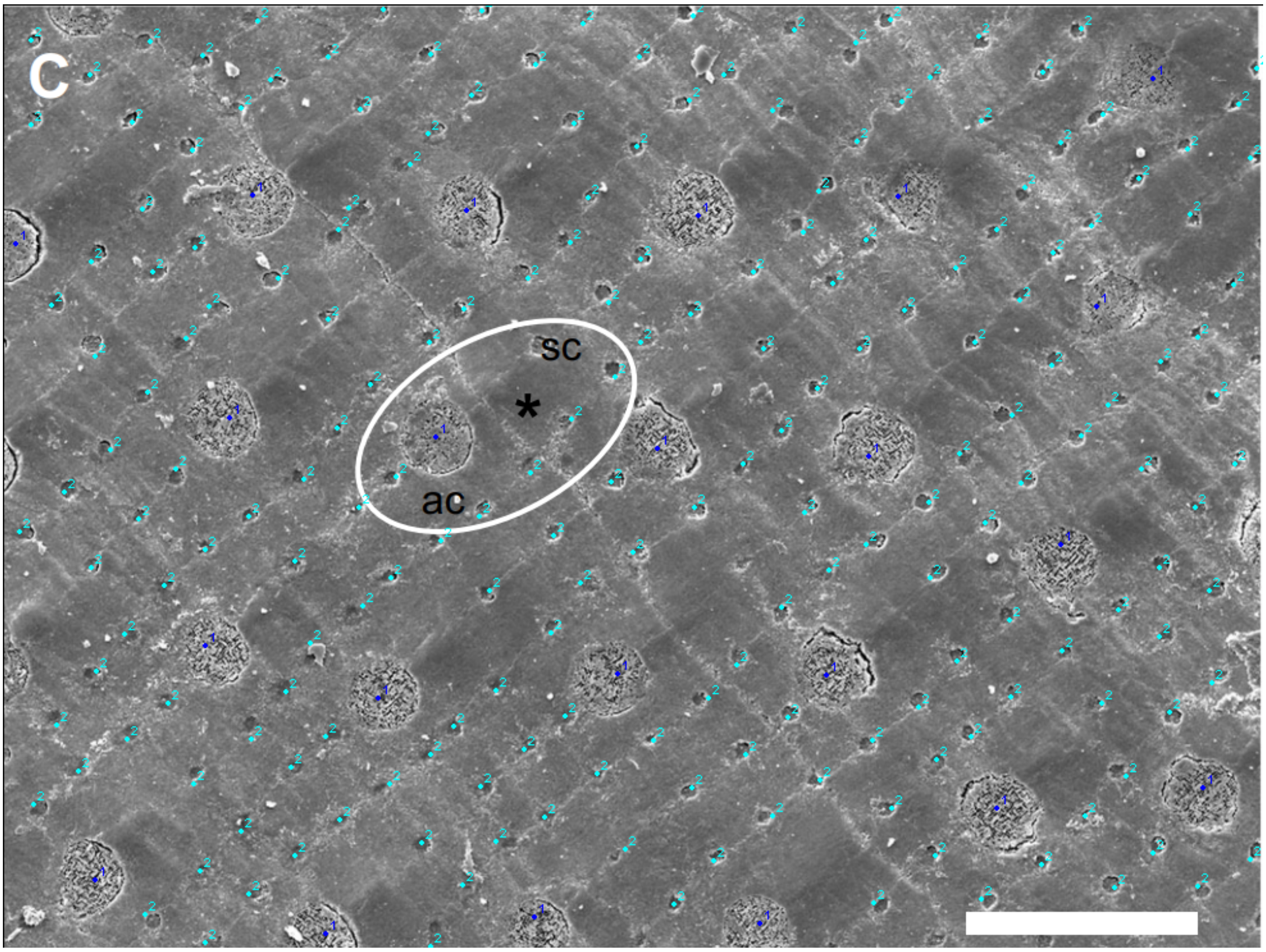

(example image from (14))

From these counts, we could calculate density in aesthetes/mm2 for each chiton by calibrating the image with the scale bar. We were limited by the availability of SEM images of sufficient magnification. The resulting aesthete data is:

| **Species** | **Density** | **Citation** | **Data From** | **Valve** |
|---|---|---|---|---|
| Acanthopleura gemmata | 2263 | Brooker 2003 | SEM | Intermediate |
| Acanthopleura granulata | 2752 | Connors 2014 | SEM, µCT | Intermediate |
| Acanthopleura spinosa | 2832 | Brooker 2003 | SEM | Intermediate |
| Callochiton achitinus | 2235 | Baxter et al. 2009 | SEM, histology | Multiple available |
| Callochiton septemvalis | 2288 | Baxter and Jones 1984 | SEM | Intermediate |
| Chiton marmoratus | 2412 | Kingston et al. 2018 | SEM | Head |
| Chiton tuberculatus | 2520 | Kingston et al. 2018 | SEM | Head |
| Cryptoplax mystica | 114 | Currie 1989 | SEM | Head |
| Ischnochiton australis | 405 | Currie 1989 | | Head |
| Ischnochiton erythronotus | 844 | Reyes-Gomez et al. 2017 | SEM | Intermediate |
| Lepidozona mertensii | 925 | Fernandez et al. 2007 | Canal casts | Intermediate |
| Mopalia acuta | 1143 | Fernandez et al. 2007 | Canal casts | Intermediate |

| | | | | |
|---|---|---|---|---|
| Mopalia muscosa | 1200 | Fernandez et al. 2007 | Canal casts | Intermediate |
| Nutallina hyadesi | 1139 | Fernandez et al. 2007 | Canal casts | table from casts |
| Onithochiton quercinus | 1542 | Currie 1989 | SEM | Head |
| Plaxiphora velata | 807 | Fernandez et al. 2007 | Canal casts | Intermediate |
| Schizochiton incisus | 3231 | Schwabe 2010, Moseley 1885 | SEM | Intermediate |
| Stenoplax bahamensis | 336 | Reyes-Gomez et al. 2017 | SEM | Intermediate |
| Stenoplax floridana | 566 | Bullock 1985 | SEM | Multiple available |
| Stenoplax limaciformis | 1148 | Bullock 1985 | SEM | Multiple available |
| Stenoplax producta | 512 | Bullock 1985 | SEM | Multiple available |
| Stenoplax purpurascences | 1166 | Bullock 1985 | SEM | Multiple available |
| Tonicella lineata | 663 | Vendrasco et al. 2008 | Canal casts | Intermediate |
| Tonicella marmorea | 535 | Connors 2014 | µCT | Intermediate |

#### 4.3 Outgroup and Available Sequences

| **Taxon** | **NCBI Accession** |
|---|---|
| Crassostrea virginica | GCA_002022765.4 |
| Chrysomallon squaminiferum | GCA_012295275.1 |
| Scutopus ventrolineatus | SRX3872460 |
| Apodomenia enigmatica | SRX3872468 |
| Proneomenia sluteri | SRX3872466 |
| Acanthochitona crinita | SRX2422912 |
| Cryptochiton stelleri | DRX136137 |
| Mopalia muscosa | SRX8144968 |
| Nuttallochiton mirandus | SRX8235049 |
| Tonicella lineata | SRX8145069 |
| Chaetopleura apiculata | SRX8235058 |
| Chiton marmoratus | SRX8235047 |
| Chiton tuberculatus | SRX8235048 |
| Hanleya hanleyi | SRX8235059 |

### 5 Maximum Likelihood Analyses

#### 5.1 Maximum Likelihood Tree

We made our initial phylogenetic tree in IQ-TREE 2 (57), using the model MFP+MERGE to implement Lanfear clustering to reduce a fully-partitioned model down to a best-fitting merged set of models and partitions.

```
#!/bin/bash -l

#SBATCH --job-name=iqtree
#SBATCH --workdir=/home/
#SBATCH --nodes=1
#SBATCH --cpus-per-task=16
#SBATCH --mem=96G
#SBATCH --time=72:00:00
#SBATCH --mail-type=ALL
#SBATCH --mail-user=

source /home/.bash_profile

iqtree -s FcC_supermatrix.fas -spp *.nex -bb 1000 -ntmax 16 -nt AUTO -m MFP+MERGE -rcluster 10
    -pre <date>
```

#### 5.2 Coalescence-based approach

To assess the impact of a coalescence based approach, we also used ASTRAL-Pro (Zhang et al. 2020) which assumes the multi-species coalescent to estimate a species tree from a high number of gene trees. We first generated individual gene trees for all loci with IQTree2:

```
# IQTree2 individual gene trees loop:
for i in *.tre; do
   iqtree -m -B 1000
done
```

```
#!/bin/bash -l

#SBATCH --job-name=astral
#SBATCH --nodes=1
#SBATCH --cpus-per-task=16
#SBATCH --mem=96G
#SBATCH --time=72:00:00
#SBATCH --mail-type=ALL
#SBATCH --mail-user=

source /home/user/.bash_profile

java -D"java.library.path=/home/user/bin/A-pro-master/ASTRAL-MP/lib" -jar
    /home/user/bin/A-pro-master/ASTRAL-MP/astral.1.1.6.jar -i loci.treefile -o astral
#java -jar /home/user/bin/A-pro-master/ASTRAL-MP/astral.1.1.6.jar -i loci.treefile
```

#### 5.3 RogueNaRok Leaf Instability Testing

To assess whether specific branches were unstable relative to the rest of the tree, we used RogueNaRok's leaf instability index (88). Investigating *Schizochiton*, the leaf stability was not lower for this taxon than for the majority of the tree. However, leaf stability and branch length indices did indicate that the sample of *Callochiton septemvalis* comprised a very long branch and very low stability, indicating that this taxon may be misidentified and/or responsible for errors in topology. Further testing, including

BLAST analyses, caused us to doubt the identity of *C. septemvalis*. Due to these concerns, and with plenty of remaining representatives of *Callochiton*, we removed this sample from subsequent analyses.

| **Taxon** | **lsDif** | **lsEnd** | **lsMax** |
|---|---|---|---|
| Lorica volvox | 0.993176 | 0.993053 | 0.994854 |
| Enlopochiton echinatus | 0.992726 | 0.99245 | 0.994517 |
| Enlopochiton niger | 0.992726 | 0.99245 | 0.994517 |
| Lucilina lamellosa | 0.992726 | 0.99245 | 0.994517 |
| Lucilina lamellosa MNHN | 0.992726 | 0.99245 | 0.994517 |
| New Genus new species Guam | 0.992726 | 0.99245 | 0.994517 |
| Onithochiton clade 11 | 0.992726 | 0.99245 | 0.994517 |
| Onithochiton neglectus | 0.992726 | 0.99245 | 0.994517 |
| Onithochiton noemiae | 0.992726 | 0.99245 | 0.994517 |
| Onithochiton quercinus | 0.992726 | 0.99245 | 0.994517 |
| Tonicia calbucensis | 0.992726 | 0.99245 | 0.994517 |
| Tonicia forbesii | 0.992726 | 0.99245 | 0.994517 |
| Tonicia lebruni | 0.992726 | 0.99245 | 0.994517 |
| Tonicia schrammi | 0.992726 | 0.99245 | 0.994517 |
| Acanthopleura gemmata SER | 0.99272 | 0.992429 | 0.994512 |
| Acanthopleura gemmata WA | 0.99272 | 0.992429 | 0.994512 |
| Acanthopleura granulata | 0.992717 | 0.992419 | 0.99451 |
| Acanthopleura planispina | 0.992717 | 0.992419 | 0.99451 |
| Squamopleura araucariana | 0.992618 | 0.99228 | 0.994436 |
| Squamopleura clade 22 | 0.992618 | 0.99228 | 0.994436 |
| Liolophura japonica | 0.992511 | 0.992121 | 0.994356 |
| Acanthopleura spinosa WA | 0.992503 | 0.992091 | 0.994349 |
| Acanthochitona crinita | 0.992275 | 0.992428 | 0.994179 |
| Apodomenia enigmatica | 0.992275 | 0.992428 | 0.994179 |
| Callochiton bouveti | 0.992275 | 0.992428 | 0.994179 |
| Callochiton crocinus | 0.992275 | 0.992428 | 0.994179 |
| Callochiton dentatus | 0.992275 | 0.992428 | 0.994179 |
| Callochiton sp. | 0.992275 | 0.992428 | 0.994179 |
| Chrysomallon squamiferum | 0.992275 | 0.992428 | 0.994179 |
| Crassostrea virginica | 0.992275 | 0.992428 | 0.994179 |
| Cryptoconchus floridanus | 0.992275 | 0.992428 | 0.994179 |
| Cyanoplax dentiens | 0.992275 | 0.992428 | 0.994179 |
| Nuttallochiton mirandus | 0.992275 | 0.992428 | 0.994179 |
| Plaxiphora albida | 0.992275 | 0.992428 | 0.994179 |
| Plaxiphora aurata | 0.992275 | 0.992428 | 0.994179 |
| Plaxiphora Fremblya matthewsi | 0.992275 | 0.992428 | 0.994179 |
| Proneomenia sluiteri | 0.992275 | 0.992428 | 0.994179 |
| Schizochiton incisus | 0.992275 | 0.992428 | 0.994179 |
| Scutopus ventrolineatus | 0.992275 | 0.992428 | 0.994179 |
| Rhyssoplax maldivensis | 0.992169 | 0.991556 | 0.994098 |
| Rhyssoplax olivaceus | 0.992169 | 0.991556 | 0.994098 |
| Chiton cumingsii | 0.992167 | 0.991556 | 0.994098 |
| Rhyssoplax calliozonas | 0.992164 | 0.991539 | 0.994094 |
| Tegulaplax hululensis | 0.992164 | 0.991539 | 0.994094 |
| Radsia barnesi | 0.992136 | 0.991469 | 0.994074 |
| Sypharochiton pelliserpentis | 0.992136 | 0.991469 | 0.994074 |
| Mopalia hindsii | 0.992109 | 0.99214 | 0.994054 |
| Mopalia muscosaA | 0.992109 | 0.99214 | 0.994054 |
| Mopalia muscosaB | 0.992109 | 0.99214 | 0.994054 |
| Callistochiton antiquusB | 0.992078 | 0.991577 | 0.994031 |
| Callistochiton shuttleworthianus | 0.992078 | 0.991577 | 0.994031 |
| Lepidozona serrata | 0.992078 | 0.991577 | 0.994031 |
| Tonicella lineata | 0.992047 | 0.991981 | 0.994008 |
| Cryptochiton stelleri | 0.991881 | 0.991772 | 0.993883 |

| | | | |
|---|---|---|---|
| Katharina tunicata | 0.991881 | 0.991772 | 0.993883 |
| Hanleya nagelfar | 0.991494 | 0.991626 | 0.993593 |
| Leptochiton asellus | 0.991494 | 0.991626 | 0.993593 |
| Leptochiton nexus | 0.991455 | 0.991498 | 0.99353 |
| Nierstraszella c.f. lineata | 0.991455 | 0.991498 | 0.99353 |
| Oldroydia percrassa | 0.991455 | 0.991498 | 0.99353 |
| Chiton marmoratus | 0.991153 | 0.990891 | 0.993337 |
| Chiton tuberculatus | 0.991153 | 0.990891 | 0.993337 |
| Chiton virgulatus | 0.991153 | 0.990891 | 0.993337 |
| Chiton viridis | 0.991153 | 0.990891 | 0.993337 |
| Leptochiton rugatus | 0.990697 | 0.990809 | 0.992995 |
| Chiton virgulatus | 0.990119 | 0.990212 | 0.992561 |
| Lepidozona willetti | 0.98966 | 0.985988 | 0.99203 |
| Lepidozona clathrata | 0.989659 | 0.985985 | 0.992029 |
| Lepidozona mertensii | 0.989659 | 0.985985 | 0.992029 |
| Lepidozona retiporosa | 0.989659 | 0.985985 | 0.992029 |
| Tripoplax trifida | 0.989636 | 0.985893 | 0.992009 |
| Callistochiton elenensisA | 0.98958 | 0.985802 | 0.991969 |
| Callistochiton elenensisB | 0.98958 | 0.985802 | 0.991969 |
| Callistochiton palmulatus | 0.98958 | 0.985802 | 0.991969 |
| Ischnochiton hakodadensis | 0.989492 | 0.985624 | 0.991905 |
| Lepidozona radians | 0.989492 | 0.985624 | 0.991905 |
| Calloplax janeirensis | 0.988863 | 0.985201 | 0.991647 |
| Calloplax vivipara | 0.988863 | 0.985201 | 0.991647 |
| Chaetopleura lurida | 0.988863 | 0.985201 | 0.991647 |
| Chaetopleura benaventei | 0.988745 | 0.985032 | 0.991558 |
| Chaetopleura peruviana | 0.988745 | 0.985032 | 0.991558 |
| Chaetopleura apiculata | 0.98839 | 0.984524 | 0.991292 |
| Ischnochiton erythronotus | 0.988371 | 0.983058 | 0.990922 |
| Ischnochiton tridentatus | 0.988371 | 0.983058 | 0.990922 |
| Stenoplax mariposa | 0.988371 | 0.983058 | 0.990922 |
| Ischnoplax sp. BC4518 | 0.988245 | 0.982885 | 0.990828 |
| Stenoplax limaciformis | 0.988245 | 0.982885 | 0.990828 |
| Stenoplax conspicua | 0.987865 | 0.982353 | 0.990542 |
| Callistochiton ashbyi | 0.987861 | 0.982142 | 0.99054 |
| Ischnochiton bouryi | 0.987861 | 0.982142 | 0.99054 |
| Ischnochiton oniscus | 0.987861 | 0.982142 | 0.99054 |
| Ischnochiton RSA | 0.987861 | 0.982142 | 0.99054 |
| Ischnochiton sirenkoi | 0.987861 | 0.982142 | 0.99054 |
| Ischnochiton yerburyi | 0.987861 | 0.982142 | 0.99054 |
| Ischnochiton haploplax lentiginosus | 0.987044 | 0.987477 | 0.990255 |
| Ischnochiton maorianus | 0.987044 | 0.987477 | 0.990255 |
| Ischnochiton stramineus | 0.987044 | 0.987477 | 0.990255 |
| Ischnochiton versicolor | 0.987044 | 0.987477 | 0.990255 |
| Tonicina zschaui | 0.987044 | 0.987477 | 0.990255 |
| Ischnochiton comptus | 0.987014 | 0.987414 | 0.990233 |
| Ischnochiton winckworthi | 0.987014 | 0.987414 | 0.990233 |
| Stenochiton cymodocealis | 0.987014 | 0.987414 | 0.990233 |
| Ischnochiton pusio | 0.986155 | 0.986254 | 0.989589 |
| Stenosemus mexicanus | 0.986155 | 0.986254 | 0.989589 |
| Loricella angasi | 0.977905 | 0.968834 | 0.983401 |
| Callochiton septemvalvis | 0.754596 | 0.791328 | 0.815919 |

### 6 Bayesian Inference Analyses/Divergence Time Estimation

#### 6.1 Dating Overview

We dated our phylogeny in MrBayes following (Zhang and Wang 2019) and (Zhang 2017) https://arxiv.org/pdf/1603.05707.pdf.

#### 6.2 SortaDate

SortaDate is an algorithm that ranks a set of gene trees by three factors: 1) consistency with the species tree, in our case the tree inferred from all genes in ASTRAL-Pro, called "bipartition"; 2) consistent root-to-tip branch lengths across the phylogeny, called "treelength", and; 3) the "clock-like" nature of each gene, called "root-to-tip". This allowed us to selected a recommended 15 loci to use for divergence time estimation. To test the impact of removing data by using only a subsection of genes, we repeated the analysis with 20 and 25 genes selected as well, but no large differences resulted in the dates generated for our phylogeny. We implemented SortaDate as follows:

First we created single-gene trees for each locus. Note that SortaDate requires a SINGLE TAXON as the root of the phylogeny, so for these analyses we removed *C. virginica* from all alignments:

```
#IQTree won't permit sequences that are empty (all Xs) so these must be removed first:
#linearize sequences
for file in *.fas; do
    awk '/^>/ {printf("%s%s\n",(N>0?"\n":""),$0);N++;next;} {printf("%s",$0);} END
        {printf("\n");}' $file > $file.lin
done
#delete any lines that contain an X
for file in *.fas.lin; do
sed '/X/d' $file > $file.sed
done
#remove resulting blank headers
for file in *.sed; do
awk 'BEGIN {RS = ">" ; FS = "\n" ; ORS = ""} {if ($2) print ">"$0}' $file > $file.awk
done
```

Switching to HPC:

```
#!/bin/bash -l

#SBATCH --job-name=iqtree_loop
#SBATCH --nodes=1
#SBATCH --cpus-per-task=16
#SBATCH --mem=96G
#SBATCH --time=72:00:00
#SBATCH --mail-type=ALL
#SBATCH --mail-user=

source /home/user/.bash_profile

for file in *.fas; do
       iqtree -s $file -bb 1000 -ntmax 16 -nt AUTO -m MFP -pre $file
done
```

Note: SortaDate requires library Armadillo; just install the version provided within the SortaDate binary, as other methods are not recognized. Next, we ran SortaDate:

```
for file in *.contree; do nw_reroot $file Csqua > $file.newick; done
python /home/user/SortaDate/src/get_var_length.py /home/user/Desktop/2021.08.23_SortaDate
    --flend .contree.newick --outf result1.txt --outg Csqua
python /home/user/SortaDate/src/get_bp_genetrees.py /home/user/Desktop/2021.08.23_SortaDate
    concat.tre --flend .contree.newick --outf result2.txt
```

```
python /home/user/SortaDate/src/combine_results.py
    /home/user/Desktop/2021.08.23_SortaDate/result1.txt
    /home/user/Desktop/2021.08.23_SortaDate/result2.txt --outf result3.txt
python /home/user/SortaDate/src/get_good_genes.py
    /home/user/Desktop/2021.08.23_SortaDate/result3.txt --max 15 --order 3,1,2 --outf
    result4.txt
```

The gene list, prioritized by the recommendations in the SortaDate manual, is:

00347.fas 01031.fas 02057.fas 03411.fas 03880.fas 04012.fas 04284.fas 04817.fas 05071.fas 05072.fas 05395.fas 05809.fas 06126.fas 09237.fas 09678.fas

#### 6.3 Fossil Constraints

Chiton fossils pose many problems, both because it is rare to find intact shell valves and also because of biases in fossilization across caldes (84). Thus, we used fossil calibrations as minimum values that allowed for uniform distributions of probabilities from the minimum to the maximum, we which define for all clades as the base of Polyplacophora itself (359 mya). Many chiton fossils are assigned to species, but these identification are often optimistic and based on morphological features that are known to vary in extant chiton groups. Thus, we applied calibrations to the best-supported taxonomic level of each fossil based on the description of characters available (e.g. a fossil identified to species may instead be used to calibrate a genus or family as 'at least X my old'.).

| Clade | Date Range | Citation(s) |
|---|---|---|
| Polyplacophora | Max 359 mya | ((*6*), (*13*)) |
| Lepidopleurida | 201.3 - 359 | ((*13*), (*5*)) |
| Chitonida | 66 - 359 | ((*2*), (*7*), (*1*), (*4*)) |
| Acanthochitonida | 33.9 - 359 | ((*7*), (*10*), (*3*)) |
| Plaxiphora | 23.03 - 359 | ((*7*), (*3*)) |
| Mopalia | 15 - 359 | ((*7*), (*3*)) |
| Tonicia | 33.9 - 359 | ((*7*)) |

#### 6.4 MrBayes Dating

Using the fossil constraints above, we ran MrBayes to date our phylogeny of chitons as follows. We stopped the run when chain discordance dropped below 0.25, higher due to the larger phylogeny. Run time was approximately 2.5 weeks.

```
#New constraints with NAMES

execute FcCsupermatrix.fas.nexus;

charset trimal.mafft.00347 = 1-455;
charset trimal.mafft.01031 = 456-677;
charset trimal.mafft.02057 = 678-1009;
charset trimal.mafft.03411 = 1010-1255;
charset trimal.mafft.03880 = 1256-1641;
charset trimal.mafft.04012 = 1642-1744;
charset trimal.mafft.04284 = 1745-1999;
charset trimal.mafft.04817 = 2000-2378;
charset trimal.mafft.05071 = 2379-2832;
charset trimal.mafft.05072 = 2833-3013;
charset trimal.mafft.05395 = 3014-3343;
charset trimal.mafft.05809 = 3344-3700;
charset trimal.mafft.06126 = 3701-4082;
charset trimal.mafft.09237 = 4083-4366;
charset trimal.mafft.09678 = 4367-4772;

partition fifteen = 15: trimal.mafft.00347, trimal.mafft.01031, trimal.mafft.02057,
    trimal.mafft.03411, trimal.mafft.03880, trimal.mafft.04012, trimal.mafft.04284,
```

```
    trimal.mafft.04817, trimal.mafft.05071, trimal.mafft.05072, trimal.mafft.05395,
    trimal.mafft.05809, trimal.mafft.06126, trimal.mafft.09237, trimal.mafft.09678;

set partition = fifteen;

prset aamodelpr=fixed(gtr);

prset clockratepr = lognorm(-7,0.6);

prset clockvarpr = igr;

prset igrvarpr = exp(10);

outgroup Cvirginica;

constraint chitonina = Cantiquus Cashbyi CelenensisA Cpalmulatus Cshuttleworthianus
    Cjaneirensis Cvivipara Cbouveti Ccrocinus Cdentatus Csp. Capiculata Cbenaventei Clurida
    Cperuviana Agemmata Agemmata2 Agranulata Aplanispina Aspinosa Cbarnesii Ccumingsii
    Cmarmoratus Ctuberculatus Cvirgulatus2 Cviridis Eechinatus Eniger Ljaponica Llamellosa
    Oneglectus Onoemiae Oquercinus Rcalliozona Rmaldivensis Rolivacea Saraucariana Sclade
    Spelliserpentis Thululensis Tcalbucensis Tforbesii Tlebruni Tschrammi Ibouryi Icomptus
    Ierythronotus Ihakodadensis Ilentiginosus Imaorianus Ioniscus Ipusio Irsa Isirenkoi
    Istramineus Itridentatus Iversicolor Iwinckworthi Iyerburyi Isp. Scymodocealis Sconspicua
    Slimaciformis Smariposa Smexicanus Lvolvox Langasi Sincisus;
constraint chitonida = Acrinita Cstelleri Cfloridanus Lclathrata Lmertensii Lradians
    Lretiporosa Lserrata Lwilletti Ttrifida Ktunicata Mhindsii Mmuscosa Mmuscosa2 Nmirandus
    Palbida Paurata Pfremblya Cdentiens Tlineata Cantiquus Cashbyi CelenensisA Cpalmulatus
    Cshuttleworthianus Cjaneirensis Cvivipara Cbouveti Ccrocinus Cdentatus Csp. Capiculata
    Cbenaventei Clurida Cperuviana Agemmata Agemmata2 Agranulata Aplanispina Aspinosa
    Cbarnesii Ccumingsii Cmarmoratus Ctuberculatus Cvirgulatus2 Cviridis Eechinatus Eniger
    Ljaponica Llamellosa Oneglectus Onoemiae Oquercinus Rcalliozona Rmaldivensis Rolivacea
    Saraucariana Sclade Spelliserpentis Thululensis Tcalbucensis Tforbesii Tlebruni Tschrammi
    Ibouryi Icomptus Ierythronotus Ihakodadensis Ilentiginosus Imaorianus Ioniscus Ipusio Irsa
    Isirenkoi Istramineus Itridentatus Iversicolor Iwinckworthi Iyerburyi Isp. Scymodocealis
    Sconspicua Slimaciformis Smariposa Smexicanus Lvolvox Langasi Sincisus;
constraint insideschiz = Cantiquus Cashbyi CelenensisA Cpalmulatus Cshuttleworthianus
    Cjaneirensis Cvivipara Cbouveti Ccrocinus Cdentatus Csp. Capiculata Cbenaventei Clurida
    Cperuviana Agemmata Agemmata2 Agranulata Aplanispina Aspinosa Cbarnesii Ccumingsii
    Cmarmoratus Ctuberculatus Cvirgulatus2 Cviridis Eechinatus Eniger Ljaponica Llamellosa
    Oneglectus Onoemiae Oquercinus Rcalliozona Rmaldivensis Rolivacea Saraucariana Sclade
    Spelliserpentis Thululensis Tcalbucensis Tforbesii Tlebruni Tschrammi Ibouryi Icomptus
    Ierythronotus Ihakodadensis Ilentiginosus Imaorianus Ioniscus Ipusio Irsa Isirenkoi
    Istramineus Itridentatus Iversicolor Iwinckworthi Iyerburyi Isp. Scymodocealis Sconspicua
    Slimaciformis Smariposa Smexicanus Lvolvox Langasi;
constraint acanthochitonina = Acrinita Cstelleri Cfloridanus Lclathrata Lmertensii Lradians
    Lretiporosa Lserrata Lwilletti Ttrifida Ktunicata Mhindsii Mmuscosa Mmuscosa2 Nmirandus
    Palbida Paurata Pfremblya Cdentiens Tlineata;
constraint tonicia = Tcalbucensis Tforbesii Tlebruni Tschrammi;
constraint plaxiphora = Palbida Paurata Pfremblya;
constraint mopalia = Mhindsii Mmuscosa Mmuscosa2;
constraint chiton = Cbarnesii Ccumingsii Cmarmoratus Ctuberculatus Cvirgulatus2 Cviridis;
constraint callochiton = Cbouveti Ccrocinus Cdentatus Csp.;
constraint rhyssoplax = Rcalliozona Rmaldivensis Rolivacea;
constraint lepidopleurina = Hhanleyi Lasellus Lnexus Lrugatus Opercrassa Nc.f.lineata;
constraint aplac = Aenigmatica Psluiteri Sventrolineatus;
constraint max = Acrinita Cstelleri Cfloridanus Lclathrata Lmertensii Lradians Lretiporosa
    Lserrata Lwilletti Ttrifida Ktunicata Mhindsii Mmuscosa Mmuscosa2 Nmirandus Palbida
    Paurata Pfremblya Cdentiens Tlineata Cantiquus Cashbyi CelenensisA Cpalmulatus
    Cshuttleworthianus Cjaneirensis Cvivipara Cbouveti Ccrocinus Cdentatus Csp. Capiculata
    Cbenaventei Clurida Cperuviana Agemmata Agemmata2 Agranulata Aplanispina Aspinosa
    Cbarnesii Ccumingsii Cmarmoratus Ctuberculatus Cvirgulatus2 Cviridis Eechinatus Eniger
    Ljaponica Llamellosa Oneglectus Onoemiae Oquercinus Rcalliozona Rmaldivensis Rolivacea
    Saraucariana Sclade Spelliserpentis Thululensis Tcalbucensis Tforbesii Tlebruni Tschrammi
```

```
    Ibouryi Icomptus Ierythronotus Ihakodadensis Ilentiginosus Imaorianus Ioniscus Ipusio Irsa
    Isirenkoi Istramineus Itridentatus Iversicolor Iwinckworthi Iyerburyi Isp. Scymodocealis
    Sconspicua Slimaciformis Smariposa Smexicanus Lvolvox Langasi Sincisus Hhanleyi Lasellus
    Lnexus Lrugatus Opercrassa Nc.f.lineata Aenigmatica Psluiteri Sventrolineatus;

calibrate tonicia = unif(33.9, 359);
calibrate max = unif(359, 549);
calibrate mopalia = unif(15, 359);
calibrate rhyssoplax = unif(41.2 , 359);
calibrate plaxiphora = unif(23.03, 359);
calibrate lepidopleurina = unif(201.3, 359);
calibrate chitonida = unif(66, 359);
calibrate acanthochitonina = unif(33.9, 359);

prset topologypr = constraints(chitonina, chitonida, insideschiz, acanthochitonina, tonicia,
    plaxiphora, mopalia, chiton, callochiton, rhyssoplax, lepidopleurina, aplac, max);
prset nodeagepr = calibrated;
prset brlenspr = clock:fossilization;
prset samplestrat = diversity;
prset sampleprob = 0.0001;
prset speciationpr = exp(10);
prset extinctionpr = beta(1,1);
prset fossilizationpr = beta(1,1);
prset treeagepr = offsetexp(425,549);

mcmcp nrun = 2 nchain = 4 ngen = 50000 samplefr = 5000;
mcmcp printfr = 5000 diagnfr = 5000;

mcmc
```

### 7 Ancestral State Reconstruction (ASR)

#### 7.1 Physiological States for ASR

Our states table was as follows:

| Taxon | Aesthetes | Eyespots | Shell Eyes |
|---|---|---|---|
| Acanthochitona fascicularis | 1 | 0 | 1 |
| Acanthopleura gemmata | 1 | 0 | 1 |
| Acanthopleura gemmata2 | 1 | 0 | 1 |
| Acanthopleura granulata | 1 | 0 | 1 |
| Acanthopleura planispina | 1 | 0 | 1 |
| Acanthopleura spinosa | 1 | 0 | 1 |
| Apodomenia enigmatica | | | |
| Callistochiton antiquus B | 1 | 0 | 0 |
| Callistochiton ashbyi | 1 | 0 | 0 |
| Callistochiton elenensis A | 1 | 0 | 0 |
| Callistochiton palmulatus | 1 | 0 | 0 |
| Callistochiton shuttleworthianus | 1 | 0 | 0 |
| Callochiton bouveti | 1 | 1 | 0 |
| Callochiton crocinus | 1 | 1 | 0 |
| Callochiton dentatus | 1 | 1 | 0 |
| Callochiton sp. | | | |
| Calloplax janeirensis | 1 | 0 | 0 |
| Calloplax vivipara | 1 | 0 | 0 |
| Chaetopleura apiculata | 1 | 0 | 0 |
| Chaetopleura benaventei | 1 | 0 | 0 |
| Chaetopleura lurida | 1 | 0 | 0 |
| Chaetopleura peruviana | 1 | 0 | 0 |
| Chiton cumingsii | 1 | 0 | 0 |
| Chiton marmoratus | 1 | 1 | 0 |
| Chiton tuberculatus | 1 | 1 | 0 |
| Chiton virgulatus | 1 | 1 | 0 |
| Chiton viridis | 1 | 1 | 0 |
| Cryptoconchus floridanus | 1 | 0 | 0 |
| Chrysomallon squamiferum | | | |
| Crassostrea virginica | | | |
| Cryptochiton stelleri | 1 | 0 | 0 |
| Cyanoplax dentiens | 1 | 0 | 0 |
| Callistochiton sp.B | | | |
| Acanthopleura echinata | 1 | 0 | 0 |
| Enoplochiton niger | 1 | 0 | 1 |
| Hanleya hanleyi | 1 | 0 | 0 |
| Ischnochiton bouryi | 1 | 0 | 0 |
| Ischnochiton comptus | 1 | 0 | 0 |
| Ischnochiton erythronotus | 1 | 0 | 0 |
| Ischnochiton hakodadensis | 1 | 0 | 0 |
| Ischnochiton lentiginosus | 1 | 0 | 0 |
| Ischnochiton maorianus | 1 | 0 | 0 |
| Ischnochiton oniscus | 1 | 0 | 0 |
| Ischnochiton pusio | 1 | 0 | 0 |
| Ischnochiton sp. | 1 | 0 | 0 |
| Ischnochiton sirenkoi | 1 | 0 | 0 |
| Ischnochiton stramineus | 1 | 0 | 0 |
| Ischnochiton tridentatus | 1 | 0 | 0 |
| Ischnochiton versicolor | 1 | 0 | 0 |
| Ischnochiton winckworthi | 1 | 0 | 0 |
| Ischnochiton yerburyi | 1 | 0 | 0 |
| Ischnoplax sp. | | | |

| | | | |
|---|---|---|---|
| Katharina tunicata | 1 | 0 | 0 |
| Lepidozona clathrata | 1 | 0 | 0 |
| Lepidozona mertensii | 1 | 0 | 0 |
| Lepidozona radians | 1 | 0 | 0 |
| Lepidozona retiporosa | 1 | 0 | 0 |
| Lepidozona serrata | 1 | 0 | 0 |
| Lepidozona willetti | 1 | 0 | 0 |
| Leptochiton asellus | 1 | 0 | 0 |
| Leptochiton nexus | 1 | 0 | 0 |
| Leptochiton rugatus | 1 | 0 | 0 |
| Liolophura japonica | 1 | 0 | 1 |
| Lorica volvox | 1 | 0 | 0 |
| Loricella angasi | 1 | 0 | 0 |
| Lucilina lamellosa | 1 | 0 | 1 |
| Mopalia muscosa2 | 1 | 0 | 0 |
| Mopalia hindsii | 1 | 0 | 0 |
| Mopalia muscosa | 1 | 0 | 0 |
| Onithochiton spGuam | | | |
| Nierstraszella cf.lineata | 1 | 0 | 0 |
| Nuttallochiton mirandus | 1 | 0 | 0 |
| Oldroydia percrassa | 1 | 0 | 0 |
| Onithochiton sp.B Indonesia | | | |
| Onithochiton neglectus | 1 | 0 | 1 |
| Onithochiton noemiae | 1 | 0 | 1 |
| Onithochiton quercinus | 1 | 0 | 1 |
| Plaxiphora albida | 1 | 0 | 0 |
| Plaxiphora aurata | 1 | 0 | 0 |
| Plaxiphora matthewsi | 1 | 0 | 0 |
| Proneomenia sluiteri | | | |
| Radsia barnesii | 1 | 0 | 0 |
| Rhyssoplax calliozona | 1 | 0 | 0 |
| Rhyssoplax maldivensis | 1 | 0 | 0 |
| Rhyssoplax olivacea | 1 | 0 | 0 |
| Schizochiton incisus | 1 | 0 | 1 |
| Squamopleura araucariana | 1 | 0 | 1 |
| Squamopleura sp. Indonesia | | | |
| Stenochiton cymodocealis | 1 | 0 | 0 |
| Stenoplax conspicua | 1 | 0 | 0 |
| Stenoplax limaciformis | 1 | 0 | 0 |
| Stenoplax mariposa | 1 | 0 | 0 |
| Stenosemus mexicanus | 1 | 0 | 0 |
| Scutopus ventrolineatus | | | |
| Sypharochiton pelliserpentis | 1 | 0 | 0 |
| Tegulaplax hululensis | 1 | 0 | 0 |
| Tonicella lineata | 1 | 0 | 0 |
| Tonicia calbucensis | 1 | 0 | 1 |
| Tonicia forbesii | 1 | 0 | 1 |
| Tonicia lebruni | 1 | 0 | 1 |
| Tonicia schrammi | 1 | 0 | 1 |
| Tonicina zschaui | 1 | 0 | 1 |
| Tripoplax trifida | 1 | 0 | 0 |

#### 7.2 Ancestral State Reconstruction

For BINARY traits (eyespots and shell eyes):
We used RayDISC, a component of corHMM ((12))in R.

```
#### RayDISC ASR w/ Rooted tree
library(ape)
library(phytools)
library(corHMM)
library(castor)
library(plyr)
library(diversitree)

#DECLARE TREE FILE NAME
treefile= "renamedconcat.contree.newick"
#READ IN TREE
chitontree <- ape::read.tree(treefile);phytools::read.newick(treefile)
#GET LIST OF TAXA
chitontree$tip.label

#DATA
filename = "ML_ASR.txt"
chitondata <- read.table("BigChitonTree_STATESslitseyespoteye.csv", header = TRUE, sep = ",",
    as.is = FALSE)

#rename your nodes in the tree; allows for annotation later in iTOL
prefix<-"node"
suffix<-seq(1:104)
chitontree$node.label[chitontree$node.label=="OROOT"]<-""
chitontree$node.label[chitontree$node.label==""]<-paste(prefix, suffix, sep= "")
#Remove NAs from node.length; causes issues in rayDISC ASR
chitontree$edge.length[is.na(chitontree$edge.length)] <- 0
write.tree(chitontree, file="bigchitontree_nodelabel")

### ASR
## Perform ancestral state estimation, using an asymmetric model of evolution and marginal
## reconstruction of ancestral states

#charnum= column it's graphing
#To decide which model is best
reconbac_ER <- rayDISC(chitontree,chitondata,model="ER", ntraits=1, charnum=1,
    node.states="marginal")
reconbac_ARD <- rayDISC(chitontree,chitondata,model="ARD", ntraits=1, charnum=1,
    node.states="marginal")
reconbac_SYM <- rayDISC(chitontree,chitondata,model="SYM", ntraits=1, charnum=1,
    node.states="marginal")

## Plot reconstructions on tree of a random model
plot <- plotRECON(chitontree,reconbac_ARD$states, title = "Chiton ASR w/ rayDISC ARD model",
    piecolors = c("darkseagreen2", "cyan3"), cex = 0.5, pie.cex = 0.3, height = 60, width =
    10, file = "ChitonASRraydisc.pdf")

write.csv(reconbac_ER$states, file="statesER-slit.csv")
write.csv(reconbac_ARD$states, file="statesARD-slit.csv")
write.csv(reconbac_SYM$states, file="statesSYM-slit.csv")

#writes data into a table so you can input into iToL to visualize
write.table(reconbac_ER$states, file="chiton_slits_statesER.txt", sep="\t")
```

For MERISTIC traits (maximum slit number), we used phytools:

```
#phytools
library(phytools)

## read tree from file
chiton.tree<-read.tree("treeforASR.newick_underscored_noquotes.tre")

## plot tree
plotTree(chiton.tree,type="fan",ftype="i")
chiton.tree$tip.label

## read data
svl<-read.csv("BigChitonTree_SLITS.csv",row.names=1)
svl<-as.matrix(svl)[,1]
svl

## Must be .tre format; you can just rename the .newick to .tre
## NO SPACES ONLY UNDERSCORES
## Check your tip labels for quotes

fit<-fastAnc(chiton.tree,svl,vars=TRUE,CI=TRUE)
fit

obj<-contMap(chiton.tree,svl,plot=FALSE)
plot(obj,type="phylogram",legend=0.7*max(nodeHeights(chiton.tree)),
    fsize=c(0.7,0.9))
```

#### 7.3 Verifying that the placement of *Schizochiton* does not impact ASR results

To test whether the predictions that chitons evolved visual systems four, separate times held, even if the placement of *Schizochiton* were to change, we artificially constrained our phylogeny to place *Schizochiton* close to genus Tonicia, a placement suggested by the highly-pectinated insertion plates.

*Schizochiton* is consistently drawn to the base of whatever clade it is constrained to within the Chitonina. We therefore concluded that if *Schizochiton* were to appear in the Chitonidae, as supported by the presence of pectinated insertion plates, the overall result that chitons evolved a visual system four distinct times would still hold because ASR indicates separate origins of shell eyes even then.

If we force a topology such that all chitons with shell eyes are monophyletic (a single origin of shell eyes), and perform topology testing in IQTree2 to assess the likelihood of the two trees, we find absolute support for the placement of *Schizochiton* at the base of Chitonina where it appears in our unconstrained tree:

| Tree | logL | deltaL | bp-RELL | p-KH | p-SH | p-WKH | p-WSH | c-ELW | p-AU |
|---|---|---|---|---|---|---|---|---|---|
| 1 | -1591193.439 | 4336.2 | 0.0000 - | 0.0000 - | 0.0000 - | 0.0000 - | 0.0000 - | 0.0000 - | 0.00229 - |

where
deltaL : logL difference from the maximal logl in the set.
bp-RELL : bootstrap proportion using RELL method (Kishino et al. 1990).
p-KH : p-value of one sided Kishino-Hasegawa test (1989).
p-SH : p-value of Shimodaira-Hasegawa test (2000).
c-ELW : Expected Likelihood Weight (Strimmer and Rambaut 2002).
p-AU : p-value of approximately unbiased (AU) test (Shimodaira, 2002).
and
Tree 1 is constrained, and is compared to our unconstrained best tree.

All topology tests reject the constrained tree because it is significantly worse than the unconstrained tree (indicated by -), so we reject the constrained tree.

The constraint tree used for analysis is: ((de3230, de6119, AGRA, de2429, de3229, de6324, de6128, de8602, IM-2013-14546, de-C17-GK, de7226, ONOE, de3369, de8006, SARA, de-C16-GK, IM-2013-14442,(de7505, de7353, TONI, de8093)), de8294);

#### 8 Phylomorphospace Construction (Figure 2)

To create a phylomorphospace of chitons using slits and aesthetes, we used phytools (90) in R:

```
#This script takes in a treefile in newick format, as well as a data frame of states (slits
    and aesthetes). Note that the phylogeny must contain only taxa with established states.
library(phytools)
#DECLARE TREE FILE NAME
treefile= "FixedSimpleChitonTree.newick"
#READ IN TREE
chitontree <- ape::read.tree(treefile);phytools::read.newick(treefile)
#GET LIST OF TAXA
chitontree$tip.label

statesrownames <- data.frame(StateMatrix_Groups, row.names = 1)
states <- as.matrix(statesrownames)

phylomorphospace(chitontree, states, A=NULL)
```

### 9 Raw Sequence Data Availability

All data files, alignments, and input/output files are available online at ¡FIGSHARE/DRYAD/etc.¿
